# Microglia drive pockets of neuroinflammation in middle age

**DOI:** 10.1101/2021.02.02.429070

**Authors:** Eric N. Moca, Daniela Lecca, Keenan T. Hope, David Tweedie, Shanaya Sidhu, Lindsay Masukawa, Hannah Sitoy, Rose Mathew, Daniel R. Saban, Nigel H. Greig, Lindsay M. De Biase

## Abstract

During aging, microglia produce inflammatory factors, show reduced tissue surveillance, altered interactions with synapses, and prolonged responses to insults, positioning these cells to have profound impact on the functional integrity of nearby neurons. We and others recently showed that microglial attributes differ significantly across brain regions and CNS insults in young adult mice. However, the degree to which microglial properties vary during aging is largely unexplored. Here, we analyze and manipulate microglial aging within the basal ganglia, brain circuits that exhibit prominent regional microglial heterogeneity and where neurons are vulnerable to functional decline and neurodegenerative disease. We demonstrate that microglia in the ventral tegmental area (VTA) and substantia nigra pars compacta (SNc) exhibit unique and premature responses to aging, compared to microglia elsewhere. This is associated with ‘pockets’ of VTA/SNc neuroinflammation that are likely to compromise local synaptic function as early as middle age. Surprisingly, these early aging responses of VTA and SNc microglia do not appear to be driven by systemic inflammation, local neuron death, or early responses of astrocytes to aging. Finally, CX3CR1 receptor knockout can exacerbate and microglial ablation/repopulation can suppress early VTA microglial aging; these manipulations have been shown to affect brain-wide microglial aging, and our data demonstrate that their impact is not uniform throughout the CNS. Our findings reveal a previously unappreciated regional variation in the onset and magnitude of microglial aging responses and suggest that there may be important links between local microglial aging and vulnerability of nearby neurons to functional decline and disease.

## INTRODUCTION

During aging, microglia produce inflammatory factors^1,2^, induce inflammatory phenotypes in astrocytes^3,4^, and exhibit prolonged injury responses^5^. Cell process motility of microglia also decreases^6,7^, suggesting that microglial-based tissue surveillance and homeostasis are compromised. Finally, signals that promote microglial phagocytosis of synapses increase with aging and they engulf synapses during pre-symptomatic neurodegeneration^8,9^. These findings argue that microglia are likely key determinants of changes in neuronal viability and synaptic function that underlie compromised cognition and movement during aging. Indeed, global manipulation of microglial phenotype during aging can improve cognition^10^ and many recently identified risk genes for neurodegenerative disease are prominently expressed by microglia^11,12^.

Due to their ubiquitous distribution and historically recognized role in immune surveillance, microglia were generally assumed to be equivalent throughout the CNS. We and others recently showed that, instead, microglia display regionally distinct phenotypes^13–18^, suggesting that tissue homeostasis, microglial injury responses, and microglial modulation of synapses may locally vary. Focusing on basal ganglia circuits, we found that VTA and SNc microglia in young adult mice show lower cell density, simpler morphology, reduced lysosome content, and distinct gene expression compared to striatal and cortical microglia^13^. Microglial cell process motility^19^, phagocytotic activity^15^, and mechanisms of synapse elimination^20^, also vary across brain regions.

Do microglia exhibit significant regional heterogeneity in their responses to aging? Given the key role that microglia play in neuroinflammation, responses to CNS insults, and synaptic interactions, prominent regional differences in microglial responses to aging would have critical implications for functional integrity and disease vulnerability of local neuron populations. Pioneering studies have shown that gray and white matter microglia exhibit distinct responses to aging^21,22^ and transcriptome studies of microglia from aging mice reveal key differences in aging-induced changes in gene expression across brain regions^14^. Single cell RNAseq and FACS-based studies of whole brain microglia during aging reveal subsets of microglia with distinct patterns of gene expression and autofluorescence during aging^23,24^, further supporting the idea of heterogeneity in aging responses of these cells. Yet despite these indications that microglial aging is not uniform throughout the CNS, the nature and magnitude of regional differences in microglial aging is largely unexplored, particularly in brain regions with disease-vulnerable populations of neurons.

## METHODS

### Transgenic mice

#### C57Bl6 wildtype mice

Wildtype mice on the C57Bl6 background were purchased from Jackson labs (Stock # 000664).

#### CX3CR1^EGFP/+^ mice

*CX3CR1*^*EGFP/EGFP*^ breeders were originally purchased from Jackson labs (Stock # 005582). In these mice, EGFP is knocked in to the CX3CR1 (fractalkine receptor) locus. These mice were maintained on a C57Bl/6 background and all mice used for experiments were heterozygous (*CX3CR1*^*EGFP/+*^) except for analysis of homozygous *CX3CR1*^*EGFP/EGFP*^ mice in **Fig. 8** and **Fig. S6**. We have shown elsewhere that EGFP expression in the basal ganglia of these mice was specific to microglial cells and 98 ± 1% of Iba1+ microglia were EGFP+ ^13^.

#### CX3CR1^CreER-ires-EYFP^; iDTR mice

Mice expressing inducible cre-recombinase under control of the CX3CR1 promoter (*CX3CR1*^*Cre-ER-ires-EYFP*^ mice) and mice expressing cre-dependent flox-stop-diphtheria toxin receptor (*iDTR* mice) were purchased from Jackson Laboratories (Stock #021160 and #007900). Mice used in fate mapping studies were F1 progeny of BALB/cJ and *CX3CR1*^*Cre-ER-ires-EYFP*^; *iDTR* mice of C57BL/6J background. Mice were housed, treated with tamoxifen, and sacrificed for tissue collection in the lab of Daniel Saban at Duke University.

#### ALDH1L1-EGFP mice

*ALDH1L1-EGFP* breeders were obtained from Mutant Mouse Resource and Research Centers (MMRRC; Stock #011015-UCD) and were originally generated by Gensat. In these BAC transgenic mice, EGFP expression is driven by the astrocyte specific aldehyde dehydrogenase 1 family member L1 promoter. All mice used for experiments were heterozygous for the transgene.

In all experiments, mice of mixed sex were used, and the number of males and females in each analysis group was balanced. Mice were housed in normal light/dark cycle and had *ad libitum* access to food and water. All experiments were performed in strict accordance with protocols approved by the Animal Care and Use Committees at UCLA, NIDA, and Duke.

### Microglial ablation

To pharmacologically ablate microglial cells from the CNS, *CX3CR1*^*EGFP/+*^ mice age 13 – 20 mo were treated with the CSF1R antagonist PLX5622 (Plexxikon, Inc.) administered through the diet (1200mg/kg chow). Mice were maintained on PLX5622 or control AIN-76A diet for 3-4 weeks and were then returned to standard vivarium chow for 3 weeks before tissue collection for immunohistochemistry. Mice that underwent microglial ablation using this approach maintained their body weight during the PLX5622 treatment and subsequent period of microglial repopulation. These mice also appeared normal during qualitative evaluation of motor coordination and exploratory behavior. To confirm microglial cell death and quantify ablation efficacy, tissue was collected from some mice after they had been administered PLX5622 via diet for 7 days.

### Tamoxifen treatments

Tamoxifen (Sigma-Aldrich Cat# T5648) was dissolved in corn oil to a stock concentration of 20 mg/ml. *CX3CR1*^*CreER-ires-EYFP*^; *iDTR* mice were given two intraperitoneal injections of tamoxifen (75 mg/kg) with one day in between injections at 2 mo of age. Mice were then euthanized at 2 weeks after the second injection or allowed to age to 20-24 mo.

### Tissue collection and immunohistochemistry

Tissue was collected from wildtype, *CX3CR1*^*EGFP/+*^, *CX3CR1*^*EGFP/EGFP*^, *ALDH1L1-EGFP*, or *CX3CR1*^*CreER-ires-EYFP*^*;iDTR* mice between of 2-24 mo of age. Mice were deeply anesthetized with an injection of Euthasol (sodium pentobarbital 150 mg/kg and sodium phenytoin 19.2 mg/kg, Virbac) and perfused transcardially with 1M phosphate buffered saline (PBS) followed by ice-cold 4% paraformaldehyde (PFA) in 1M PBS. Microglia show circadian-based changes in morphology ^25^; all perfusions for this study were hence performed between 9:00am-12:00pm. Brain tissue was isolated and postfixed in this solution for 4 hours at 4°C and then stored in 1M PBS with 0.1% NaAz.

For analysis of neuron number (**Fig. 7**), brains were cryoprotected in 30% sucrose, frozen, and coronal brain sections were prepared on a cryostat at a thickness of 25μm. Brain sections were collected in series, resulting in 3 sets of serially collected sections per mouse. One set of serially collected sections per mouse was stained and 3 sections at standardized anterior-posterior locations were analyzed (**Fig. S5**). For all other analyses, brains were not cryoprotected and coronal brain sections (60μm thick) were prepared on a vibratome in chilled 1M PBS. Brain sections were grouped according to well-defined anatomical parameters and for analysis of microglial cell density (**Fig. 1**,**5**,**6**,**8** and **Fig. S6**), microglial fate mapping with DTR (**Fig. 2** and **Fig. S1**), microglial cell morphology (**Fig. 3**,**6** and **Fig. S3**), and astrocyte density (**Fig. 7**), three brain sections containing nucleus accumbens (NAc) and three brain sections containing VTA were analyzed from each mouse. Brain sections were chosen using well-defined anatomical parameters and were matched for anterior-posterior location.

**Figure 1.**
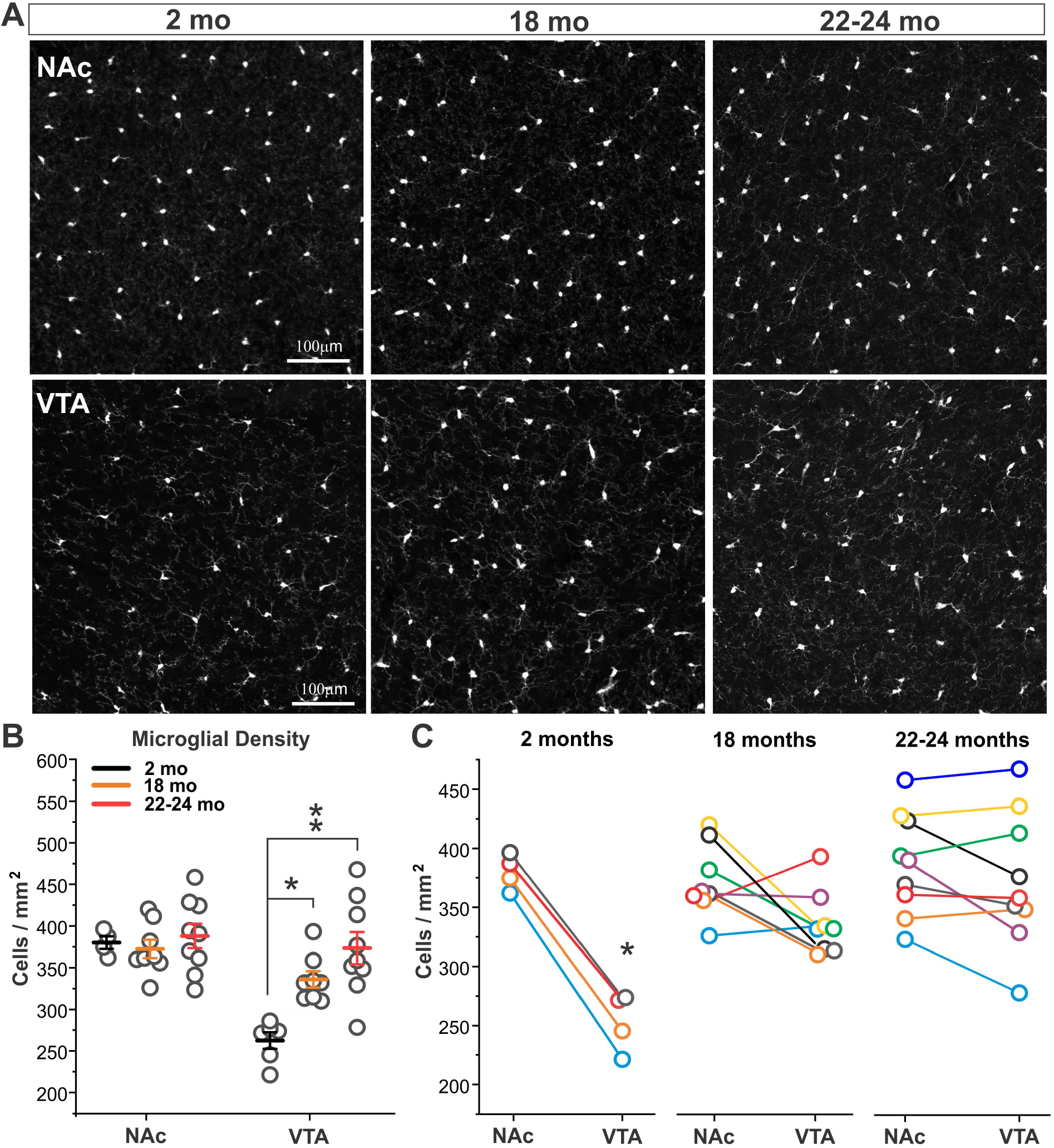
VTA microglia show early increases in cell density. **A, B** – Images and quantification of NAc and VTA microglial density in 2 month (mo), 18mo, and 22-24mo old *CX3CR1*^*EGFP/+*^ mice. Two-way ANOVA: main effect of age F(2,43) = 7.4, P = 0.002; main effect of brain region F(1,43) = 21.0, P < 0.0001; interaction F(2,43) = 5.6, P < 0.007. *P = 0.02 18mo VTA vs. 2mo VTA. ** P < 0.0001 22-24mo VTA vs. 2mo VTA. N = 4-9 mice analyzed per age. **C** – Comparison of NAc and VTA microglial density within individual mice at different ages. Data points from the same mouse colored alike. * P = 0.0002, 2mo VTA vs. NAc; 18mo VTA vs. NAc, P = 0.06 n.s.; 22-24mo VTA vs. NAc, P = 0.2 n.s., two tailed paired t-test.

**Figure 2.**
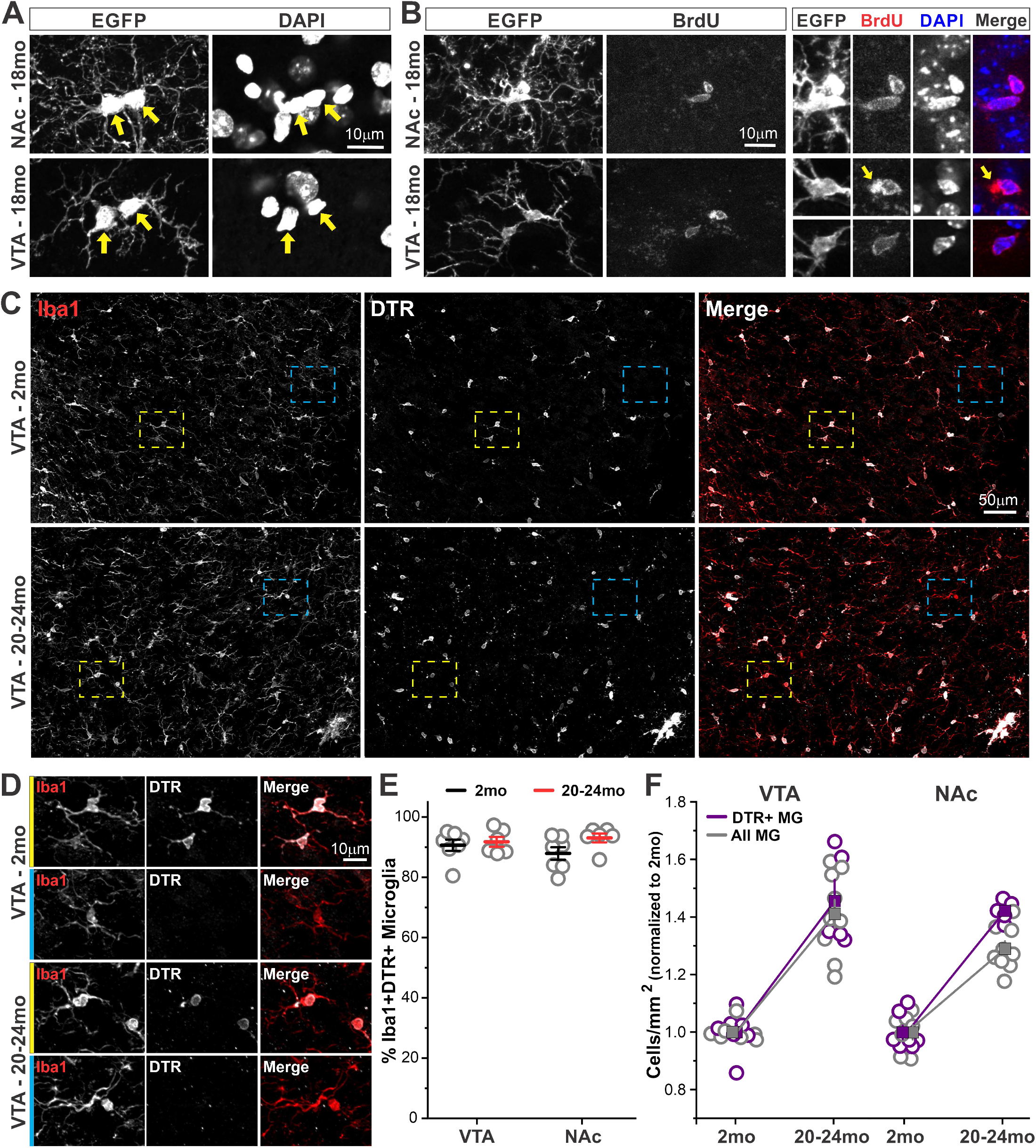
Increases in microglial density during aging arise from proliferation of resident microglia. **A** – Microglia exhibiting morphological profiles of cell division during aging. *Yellow arrows* highlight closely approximated or interconnected microglial somas with distinct DAPI+ nuclei. **B** – Examples of BrdU+ microglia during aging. *Yellow arrows* highlight lipofuscin autofluorescence that is easily distinguished from BrdU labeling of cell nuclei. **C** – Images from VTA of 2mo and 20-24mo *CX3CR1*^*CreER-IRES-EYFP*^*;iDTR* mice treated with tamoxifen at 2mo of age to label microglia present in the CNS at young adult ages. **D** – Examples of Iba1+DTR+ microglia (*yellow boxes*) and Iba1+DTR-microglia (*cyan boxes*). **E** –% of Iba1+ microglia that are also DTR+. Two-way ANOVA: main effect of age F(1,25) = 2.9, P = 0.1 n.s.; main effect of brain region F(1,25) = 0.16, P = 0.69 n.s.; interaction F(1,25) = 1.2, P = 0.28 n.s. **F** – Density increases in Iba1+DTR+ microglia compared to all (DTR+ and DTR-) microglia. Values normalized to those observed at 2mo. Two-way ANOVA: main effect of region F(1,25) = 2.7, P = 0.11 n.s.; main effect of cell population F(1,25) = 3.6, P = 0.07 n.s.; interaction F(1,25) = 0.89, P = 0.35 n.s. N = 5-9 mice analyzed per age.

**Figure 3.**
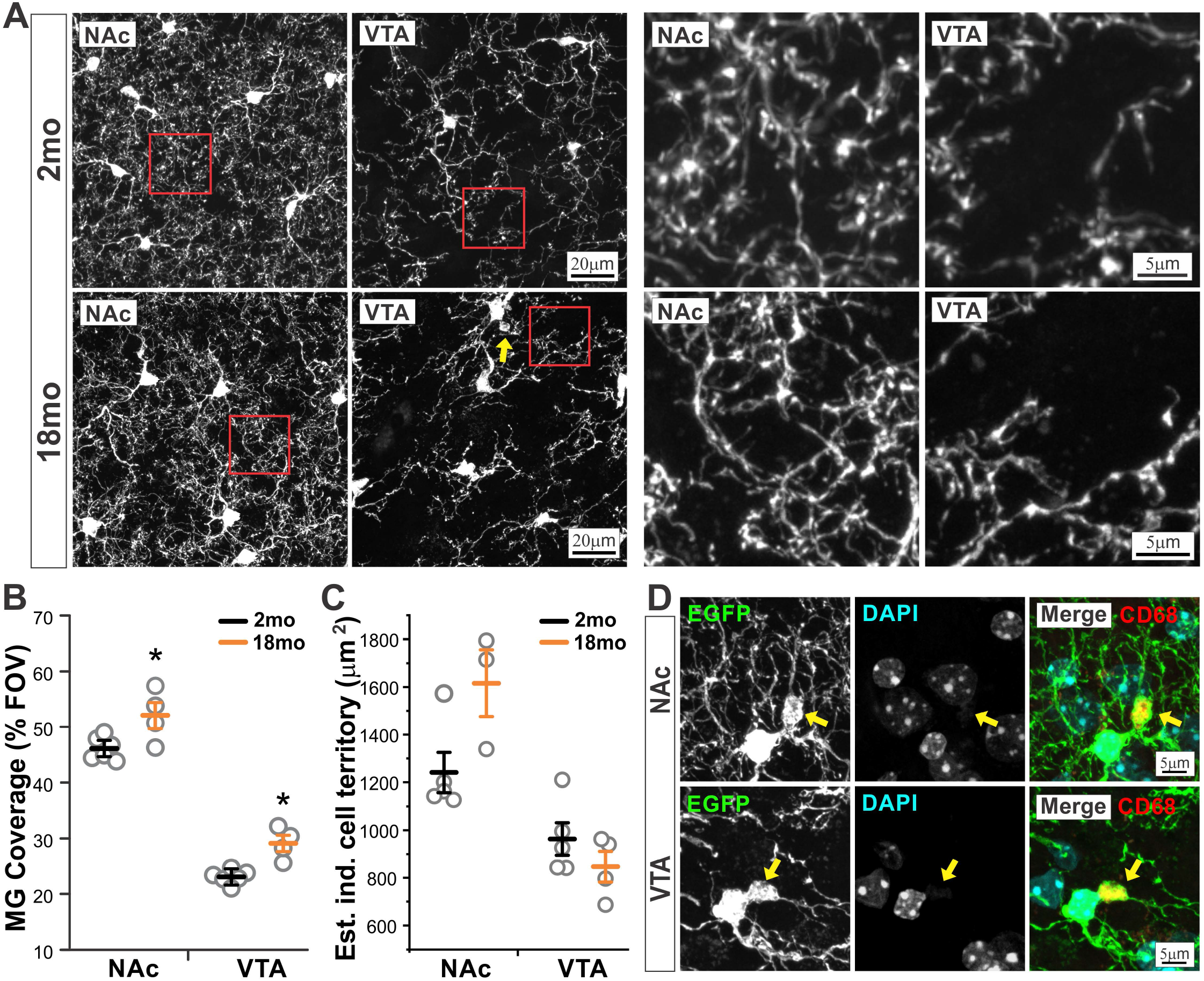
Microglial tissue coverage increases during aging. **A** – NAc and VTA microglial branching structure in 2mo and 18mo old *CX3CR1*^*EGFP/+*^ mice. Distal processes (*red boxes*) shown at higher magnification *at right. Yellow arrow* highlights an example of somatic swelling observed in some microglia at 18mo. **B** – Percent coverage of the field of view (FOV) by microglial somas and processes. Two-way ANOVA: main effect of age F(1,19) = 23.4, P = 0.0002; main effect of brain region F(1,19) = 338.6, P < 0.0001; interaction F(1,19) = 0.004, P = 0.95 n.s. *P = 0.02 18mo VTA vs. 2mo VTA, 18mo NAc vs. 2mo NAc. **C** – Estimated tissue coverage of individual microglia (total microglial coverage in FOV / # of microglia present in FOV). **D** – Examples of somatic swellings (*yellows arrows*) from VTA and NAc microglia of 18mo mice. N = 4-6 mice analyzed per age.

For immunohistochemistry, free-floating brain sections were permeabilized and blocked in 0.3% TritonX-100 and 5% Normal Donkey Serum (NDS) in 1 M PBS for 2 hours with rotation at room temperature (RT). Sections were incubated with primary antibodies prepared in 0.05% TritonX-100 and 5% NDS in 1M PBS at 4C overnight. Sections were incubated with secondary antibodies prepared in 5% NDS in 1M PBS for 2 hours at RT. Control sections incubated with secondary antibody alone did not result in labeling of cells. Primary antibodies used included the following: chicken anti-GFP (1:1000; Aves, Tigard, OR Cat#GFP-1020), goat anti-GFP (1:1000; Frontier Institute, Hokkaido, Japan, Cat#GFP-Go-Af1480), rabbit anti-Iba1 (1:500; Wako, Richmond, VA, Cat#019-19741), goat anti-DTR (R&D Systems AF259), rat anti-CD68 (1:200; clone FA-11, AbD Serotec, Raleigh, NC, Cat#MCA1957), mouse anti-tyrosine hydroxylase (1: 5000; Sigma, St. Louis, MO, Cat#T1299), chicken anti-tyrosine hydroxylase (1:500; Aves, Tigard, OR, Cat#TYH), mouse anti-NeuN (clone A60, 1:500; Millipore, Billerica, MA Cat#MAB377), rat anti-BrdU (1:500; Accurate Chemical and Scientific, Westbury, NY). Primary antibodies have been validated for use in immunohistochemistry in mouse tissue in published literature and on the manufacturer’s websites. Secondary antibodies used included the following: Alexa Fluor 488-, 594-, or 647-conjugated secondary antibodies to rabbit, mouse, goat, chicken, rat or guinea pig (1:1000; all raised in donkey; Jackson ImmunoResearch). A 1:4000 dilution of 4’,6-diamidino-2-phenylindole (DAPI) in 1M PBS was used to stain nuclei.

### Image acquisition and analysis

Fixed tissue was imaged using an Olympus FV1000 confocal microscope, a Zeiss LSM880 confocal microscope, or a Zeiss Apotome microscope. Within the NAc, analyzed images were acquired at the boundary between core and shell (identified anatomically), and include both subregions. In the VTA, analyzed images were medial to the medial lemniscus and included the parabrachial pigmented area, as well as portions of the parafasciculus retroflexus area and paranigral nucleus. For quantification of microglial or astrocyte density, stacks of confocal images (z-stacks) with a z-interval of 1.5 μm were taken through a 20x objective and imported into ImageJ software for analysis. Within Image J software, maximum intensity projections of confocal z-stacks were created and cells were counted manually. For quantification of microglial tissue coverage, z-stacks were taken with a 63x objective and a z-interval of 1.5 μm. Within ImageJ software, the mean pixel intensity of the dimmest cell processes was measured at 10-15 locations diagonally across each image. The average of these values was taken as a threshold for determining the % of pixels above (representing microglial cell somas and processes) and below (representing background) this threshold. For quantification of neuronal cell loss, tiled z-stacks of confocal images were taken with a 20x objective and a z-interval of 1.5 μm such that the entire VTA and SNc were visualized. Stitched, maximum intensity projections were imported into ImageJ and cells were counted manually. For all image analyses 3 images from separate brain sections were analyzed per mouse to obtain an average value for that mouse. For quantification of neuronal cell loss, the total number of cells across 3 stereologically matched sections was also calculated. 3-6 mice were analyzed per brain region, per age. Sample sizes were selected to be in the upper range of published immunohistochemistry experiments. Experimenters were not blind to source brain region during image analysis.

### Microdissection

*CX3CR1* ^*EGFP/+*^ mice were anesthetized with Euthasol (sodium pentobarbital 150 mg/kg and sodium phenytoin 19.2 mg/kg, Virbac) and perfused transcardially with 10 mL of oxygenated, icecold N-methyl-d-glucamine (NMDG)-based solution containing the following (in mM): 92 NMDG, 20 Hepes, 30 NaHCO3, 1.2 NaH2PO4, 2.5 KCl, 5 sodium ascorbate, 3 sodium pyruvate, 2 thiourea, 10 MgSO4, and 0.5 CaCl2, 10 glucose, pH 7.4 (310 mOsm). Brains were then rapidly dissected free. For RT-PCR and high sensitivity ELISA of tissue levels of inflammatory factors, horizontal midbrain sections (230 μm thick) and coronal forebrain sections (300 μm thick) were prepared using a vibratome in ice-cold NMDG-based cutting solution bubbled continuously with 95% O2/5% CO2. After sectioning, slices remained in ice-cold, oxygenated NMDG solution and were transferred one at a time to a glass dissecting surface under a stereoscope maintained at 4°C. The VTA was microdissected from horizontal midbrain sections, and NAc and overlying medial prefrontal cortex (mPFC) were microdissected from coronal forebrain sections using fine tipped forceps. Microdissected tissue was minced using a scalpel under the stereoscope before being transferred to eppendorf tubes containing 1 mL NMDG stored on ice. Samples were then centrifuged, NMDG removed and samples were either flash frozen (for high sensitivity ELISA protein analysis) or were resuspended in 100 μL PicoPure RNA extraction buffer and incubated at 42°C for 30 min and stored in RNase-free tubes at −80°C until further processing (for qPCR analysis). For qPCR of FACS-isolated microglia, following NMDG perfusion, rough coronal brain sections were prepared using a brain block and processed as described below.

### Tissue dissociation and flow cytometry

The striatum and overlying cortex were microdissected from anterior brain sections and the midbrain was dissected from posterior sections. Microdissected samples were minced on a glass surface maintained at 4°C. Samples were then transferred to Eppendorf tubes containing 1mL Hibernate A and were gently dissociated using sequential trituration with fire-polished glass pipettes with openings of decreasing diameter (final pipette ∼ 0.4 mm diameter opening). Resulting cell suspensions were spun down, resuspended in 300 μL 1M PBS and filtered through a 40 μm mesh filter. Throughout the experiment, samples were kept at 4°C on ice. Samples were sorted using a FACS Aria I cell sorter (BD Biosciences). The population of cells containing microglia could be readily identified based on forward scattering (FSC) and side scattering (SSC) properties. A gating strategy based on FSC and SSC width and height was used to select only single cells, and reduce debris, doublets, and dead cells. Microglial cells within this population were then identified and sorted according to EGFP expression as described previously ^13^. Microglia were FACS sorted directly into eppendorf tubes containing 50 μL PicoPure RNA extraction buffer and, following the sort, samples were incubated at 42°C for 30 min, and stored in RNase-free tubes at −80°C until further processing.

### RNA extraction and RT-PCR / ddPCR

RNA from microdissected tissues and sorted cells was isolated using the PicoPure RNA isolation kit (Arcturus Bioscience). Column filtration, washing, and elution of RNA from the columns was performed according to manufacturer’s instructions in section C of the PicoPure RNA isolation protocol. RNA concentration and integrity was then analyzed via NanoDrop spectrophotometer. Single strand cDNAs were synthesized with Superscript III first strand cDNA synthesis kit (Invitrogen, Life Technologies), according to the manufacturer’s protocol. For qPCR analysis of tissue levels of inflammatory factors, duplex RT-PCR assays were performed on technical duplicates using a FAM-labeled probe for each target gene, and a VIC-labeled probe for the endogenous control gene (*Gapdh*), along with TaqMan Advanced Fast PCR Master Mix (Cat# 4444963; Life Technologies). To avoid amplification of genomic DNA contamination, primers and probes that amplify across target gene exon–exon junctions were selected when possible. RT-PCR reactions were run in a 7500 Fast TaqMan instrument using the program: 95°C hold for 20 s, followed by 40 cycles of 95°C denaturation for 3 s, and 60°C annealing and extension for 30 s. Calculations of relative expression from Ct data were carried out according to User Bulletin #2 for ABI Prism 7900 Sequence Detection System. For each target gene, the average Ct value for the endogenous control (*Gapdh*) was subtracted from the average Ct value for the target gene, to obtain ΔCt. The relative expression was then plotted as 2^-ΔCt^. For ddPCR analysis of tissue levels of inflammatory factors, duplex assays were performed on technical duplicates using a FAM-labeled probe for each target gene, and a VIC-labeled probe for the endogenous control gene (*Gapdh*). Reactions were run in a Bio-Rad QX-200 ddPCR system (Automated Droplet Generator, Thermocycler, and Droplet Reader). Thermocycler conditions were as follows: 95°C hold for 10 min, followed by 40 cycles of 95°C denaturation for 30 s and 60°C annealing and extension for 1 min, with a final 98°C for 30 s and using ddPCR Supermix for Probes (No dUTP). Calculations of absolute expression were made within Quantasoft software according to manufacturer’s recommendations.

### Protein extraction and MesoScale analysis of cytokines

Serum samples and brain tissue samples were evaluated in duplicate using the V-PLEX Proinflammatory Panel 1 Mouse Kit (Cat # K15048D, MESO SCALE DIAGNOSTICS, LLC), and the following cytokines were assayed in each sample: Interferon-gamma (IFN-γ), Interlukin-1 beta (IL-1β), Interlukin-2 (IL-2), Interlukin-4 (IL-4), Interlukin-5 (IL-5), Interlukin-6 (IL-6), Interlukin-10 (IL-10), Interlukin-12p70 (IL-12p70), Chemokine Ligand 1 (CXCL1 or KC/GRO), Tumor necrosis factor-alpha (TNFα). All samples were assessed following the recommended protocols suggested by the manufacturer. Tissue samples (NAc: 3.9±0.9 mg (mean±sd for all tissues) (2-3 months), 4.7±2.2 mg (17-18 months), 3.1±0.5 mg (22-24 months); VTA: 2.2±1.2 mg (2-3 months), 2.7±2. mg (17-18 months), 2.1±0.7 mg (22-24 months)) were homogenized in small volume tubes, (Tissue homogenizing CKMix_WP tubes) using a Precellys Evolution plus Cryolys Evolution system (Bertin Instruments). The tissues were homogenized in approximately 65 µl of in a Tris-based lysis buffer (MSD Tris Lysis Buffer, Cat # R60TX-3) with protease and phosphatase inhibitors (Halt protease & phosphatase inhibitor single-use cocktail; Thermo Scientific, Cat # 78442). Following homogenization, lysate was extracted from the tube, placed in a fresh eppendorf tube and centrifuged (10,000 g for 10 minutes (4°C). Thereafter, the supernatant was collected into new tubes. Protein concentrations were then measured by bicinchoninic acid (BCA) assay (Micro BCA™ Protein Assay Kit, Cat # 23235; Thermo Scientific). The resulting supernatant yielded low volumes (30-40 µl of supernatant) with varying quantities of total protein, for NAc 0.6 to 5.7 µg/µl and for VTA 0.23 to 1.4 µg/µl. Serum samples were diluted 1 in 2 with the kit dilution buffer, and then loaded into the wells of a 96 well multiplex ELISA plate. Cytokine detection signals generated for each sample were compared to a cytokine standard curve, and the serum raw data levels were then calculated by use of DISCOVERY WORKBENCH (MESO SCALE DIAGNOSTICS, LLC.) and expressed as pg/ml. Data are shown as a normalized ratio relative to 2-3 month old animals. Tissue samples were mixed with the kit dilution buffer and loaded into the wells of the 96 well multiplex ELISA plate. Due to the low volumes and total protein levels recovered from the samples, samples were loaded into the wells of the ELISAs at different total protein contents. Most NAc samples were loaded at 30 µg per well, whereas the loading for NAc samples varied from 15 to 25 µg per well, depending on the sample. All VTA samples were loaded at 10 µg per well. Cytokine levels in tissues were calculated in the same way as the serum samples.

### Statistical comparisons

All graphed values are shown as mean ± SEM. Statistical details of experiments (statistical tests used, exact value of n, what n represents) can be found in the results and figures legends. In general, statistical significance was assessed using one- or two-way ANOVA (Figures 1B, 2E-F, 3B-C, 4B, 4D, 5A-B, 5D, 6A-C, 6F, 7B, 7E-F, 8C-E, 8G-I, S1B-C, S2A-D, S4A-I, S6E, S6H). Posthoc comparisons were carried out using student’s t-test with Bonferroni correction for multiple comparisons. Data are sufficiently normal and variance within groups is sufficiently similar for use of parametric tests. Paired t-tests were used to compare trends across two different regions within individual mice (Figures 1C, 4C, 5E, S6C-D, S6G).

**Figure 4.**
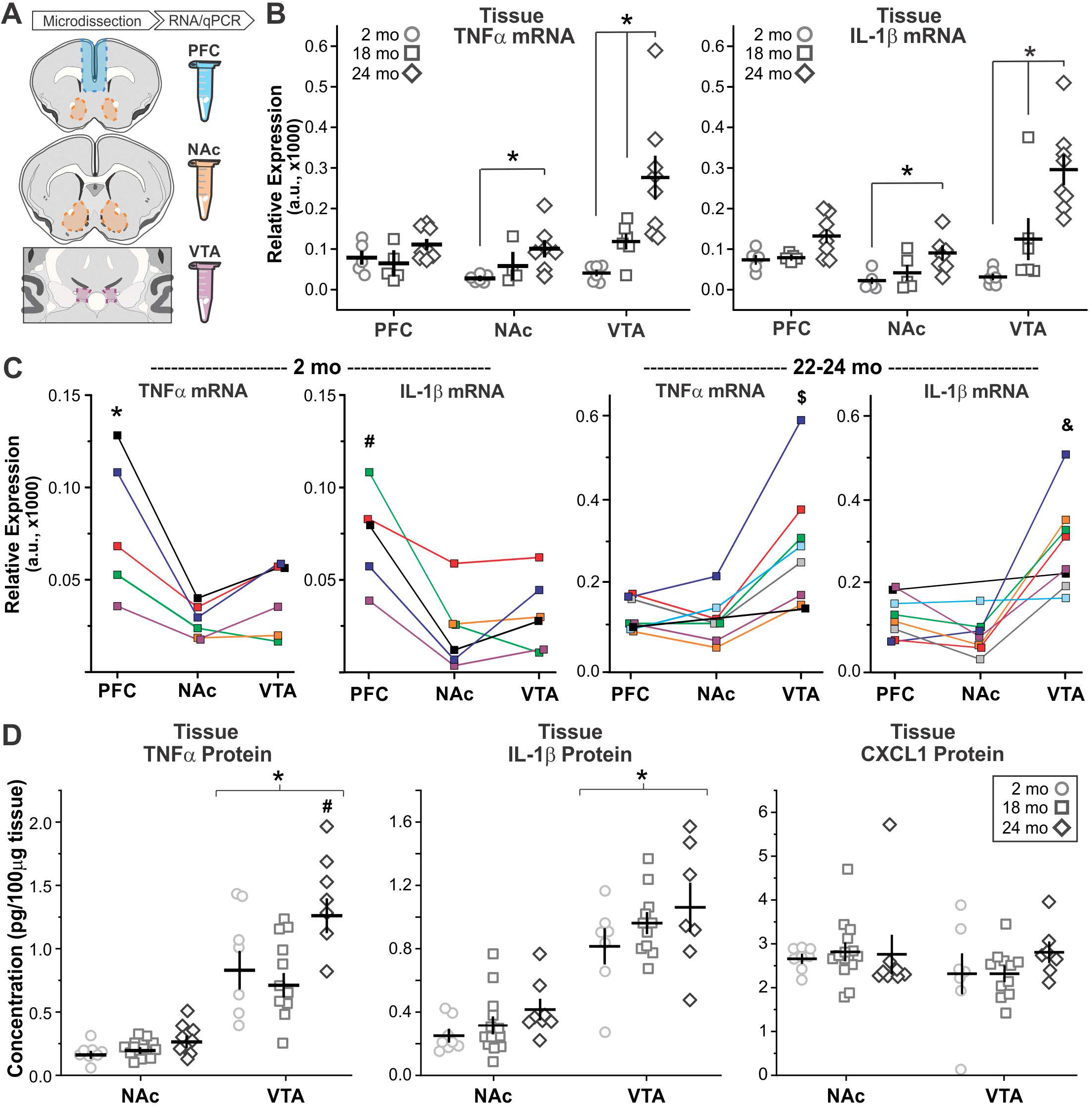
Local increases in microglial density are associated with “pockets” of inflammation. **A** – Cartoon indicating regions microdissected for mRNA and protein-level analysis in the prefrontal cortex (PFC), NAc, and VTA. **B** – Relative expression of tumor necrosis factor alpha (TNFα, *left*) and interleukin 1 beta (IL-1β, *right*) mRNA in the PFC, NAc, and VTA at 2, 18, and 24 months (mo).Two-way ANOVAS: **TNFα** main effect of age F(2,54) = 14.1, P < 0.0001; main effect of brain region F(2,54) = 6.9, P = 0.002; interaction F(4,54) = 4.0, P = 0.007; **IL-1β** main effect of age F(2,56) = 8.4, P = 0.0007; main effect of brain region F(2,56) = 5.2, P = 0.009; interaction F(4,56) = 3.3, P = 0.02. *P < 0.03. N = 5-8 mice analyzed per age, per brain region. **C** – Comparison of TNFα and IL-1β transcript across brain regions in individual mice. For each age, data points from the same mouse are colored alike. Significant paired T-tests: * P < 0.02, PFC vs. NAc and PFC vs. VTA; # P <0.02 PFC vs. NAc and PFC vs. VTA; $ P < 0.008, VTA vs. NAc and VTA vs. PFC; & P < 0.02, VTA vs. NAc and VTA vs. PFC. **D** – Local protein levels of TNFα (*left*), IL-1β (*middle*), and CXCL1 (*right*) as measured by high-sensitivity ELISA. Two-way ANOVAS **TNFα** main effect of age F(2,53) = 8.8, P = 0.0005; main effect of brain region F(1,53) = 126.9, P < 0.0001; interaction F(2,53) = 5.0, P = 0.01.; **IL-1β** main effect of age F(2,54) = 3.0, P = 0.06 n.s.; main effect of brain region F(1,54) = 95.1, P < 0.0001; interaction F(2,54) = 0.2, P = 0.8 n.s. **CXCL1**: main effect of age F(2,54) = 0.6, P = 0.56 n.s.; main effect of brain region F(1,54) = 1.5, P = 0.23 n.s.; interaction F(2,54) = 0.6, P = 0.0.57 n.s. * P < 0.0001 VTA vs. NAc. # P < 0.02 VTA 24mo vs VTA 18mo and VTA 24mo vs VTA 2mo. N = 7-13 mice analyzed per age, per brain region.

**Figure 5.**
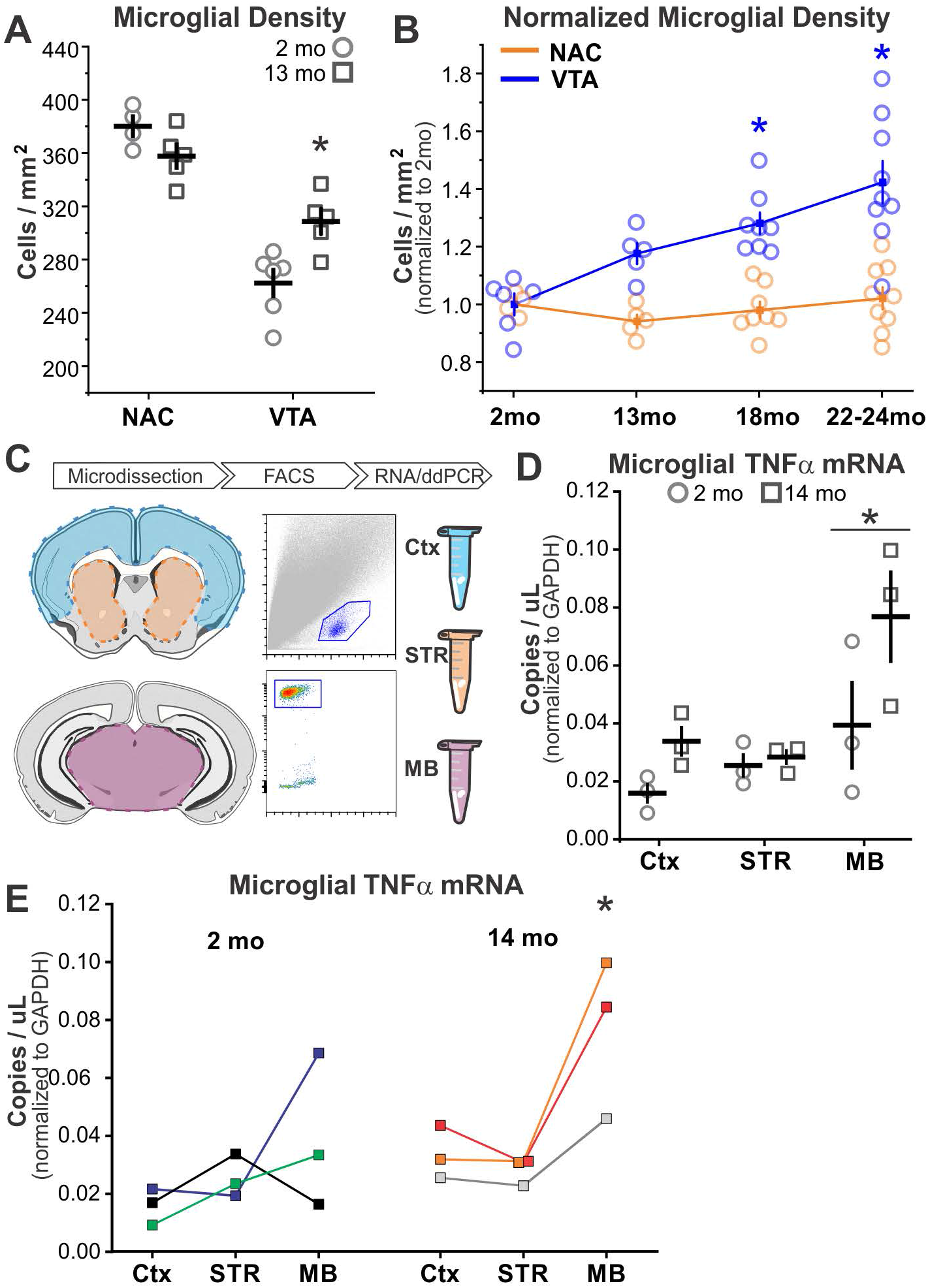
Local increases in VTA microglial density and inflammation begin as early as middle age. **A** – NAc and VTA microglial density in 2 and 13mo old *CX3CR1*^*EGFP /+*^ mice. Two-way ANOVA: main effect of age F(1,19) = 1.6, P = 0.22 n.s.; main effect of brain region F(1,19) = 77.7, P < 0.0001; interaction F(1,19) = 13.2, P = 0.002. *P = 0.01. N = 4-6 mice analyzed per age. **B** – NAc and VTA microglial density during aging normalized to values observed in 2mo animals. Two-way ANOVA: main effect of age F(3,53) = 7.9, P = 0.0002; main effect of brain region F(1,53) = 44.5, P < 0.0001; interaction F(3,53) = 5.7, P = 0.002. *P < 0.0005, 18mo VTA vs. 18mo NAc and 22-24mo VTA vs. 22-24mo NAc. **C** – Workflow schematic; midbrain (MB), striatum (STR), and cortex (CTX) were dissected from acute coronal forebrain sections followed by FACS isolation of EGFP+ microglia, RNA extraction, and ddPCR analysis. **D** – Microglial expression of TNFα transcript. Two-way ANOVA: main effect of age F(1,17) = 6.1, P = 0.03; main effect of brain region F(2,17) = 7.5, P = 0.008; interaction F(2,17) = 1.6, P = 0.24 n.s. * P < 0.03 MB vs. Ctx and MB vs. STR. **E** – Comparison of NAc and VTA microglial TNFα expression within individual mice at 2 and 14mo. Data points from same mouse are colored alike. * P < 0.05, one tailed paired t-test.

**Figure 6.**
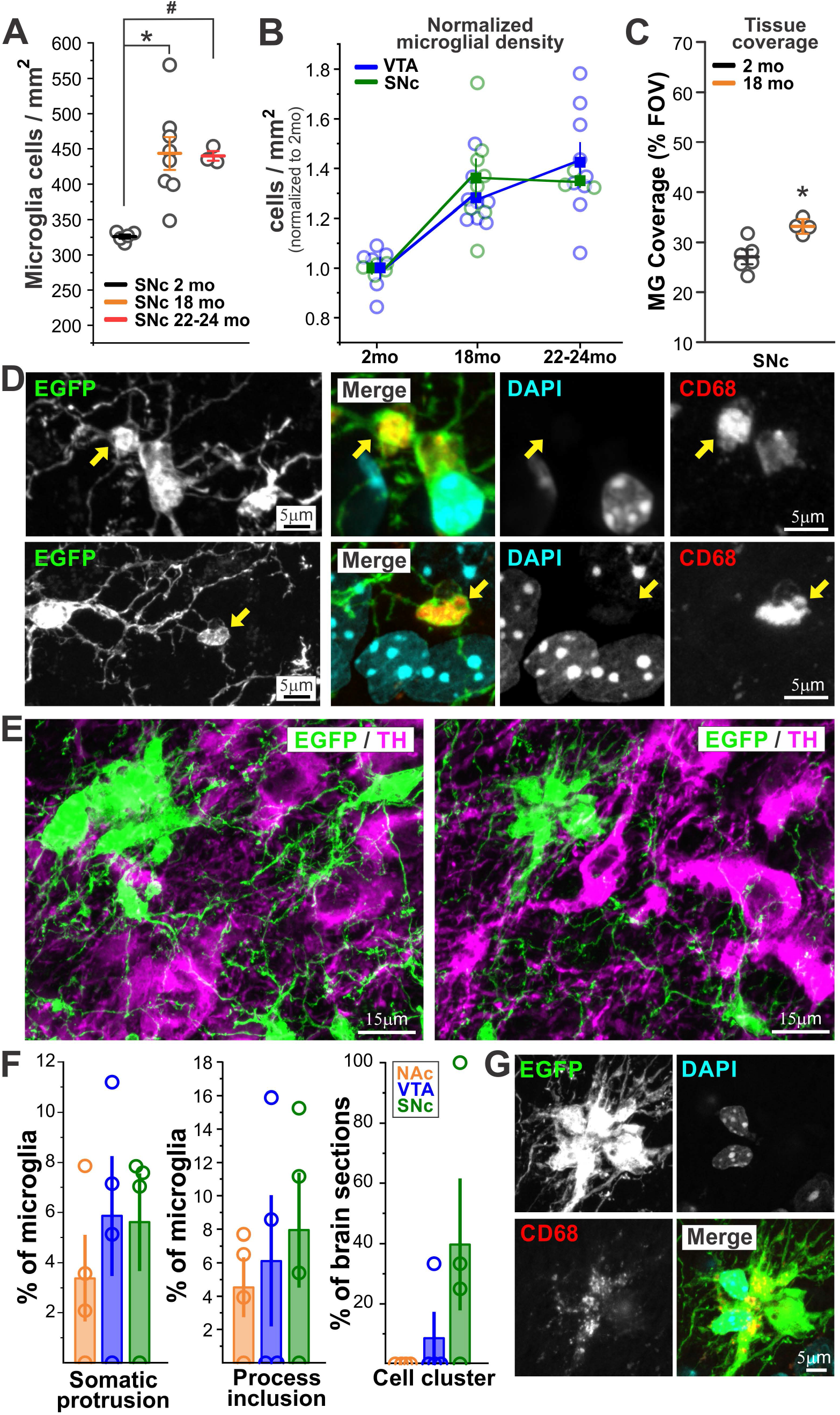
SNc microglia resemble VTA microglia during aging but also exhibit microglial clusters. **A** – SNc microglial density in 2, 18, and 22-24mo old *CX3CR1*^*EGFP /+*^ mice. One-way ANOVA F(2,15) = 9.9, P = 0.002. *P < 0.003, 18mo SNc vs. 2mo SNc. # P = 0.02 22-24mo SNc vs. 2mo SNc. **B** – Comparison of VTA and SNc microglial density increases during. Densities are normalized to values in each region at 2mo of age. Two-way ANOVA: main effect of age F(2,38) = 19.1, P < 0.0001; main effect of brain region F(1,38) = 0.001, P = 0.97 n.s.; interaction F(2,38) = 0.7, P = 0.49. **C** – Percent coverage of field of view (FOV) by microglial somas and processes. *P = 0.005, two sample t-test. **D** – Examples of somatic swellings (*top panels*) and process swellings (*bottom panels*) from SNc microglia in 18mo *CX3CR1*^*EGFP /+*^ mice. *Yellow arrows* and co-staining with DAPI and CD68 indicate that swellings contain lysosomes but not pyknotic nuclei. **E** – Examples of microglial clusters observed in SNc of 18mo old *CX3CR1*^*EGFP/+*^ mice. **F** –Frequency of microglial morphological abnormalities across the NAc, VTA, and SNc of 18mo old *CX3CR1*^*EGFP /+*^ mice. One-way ANOVAS: Somatic swellings F(2,11) = 0.5, P = 0.6 n.s.; Cell process swellings F(2,11) = 0.3, P = 0.7 n.s.; Microglial clusters F(2,11) = 2.5, P = 0.14 n.s. **G** – Co-staining of cluster *at right* in *E* with DAPI and CD68.

**Figure 7.**
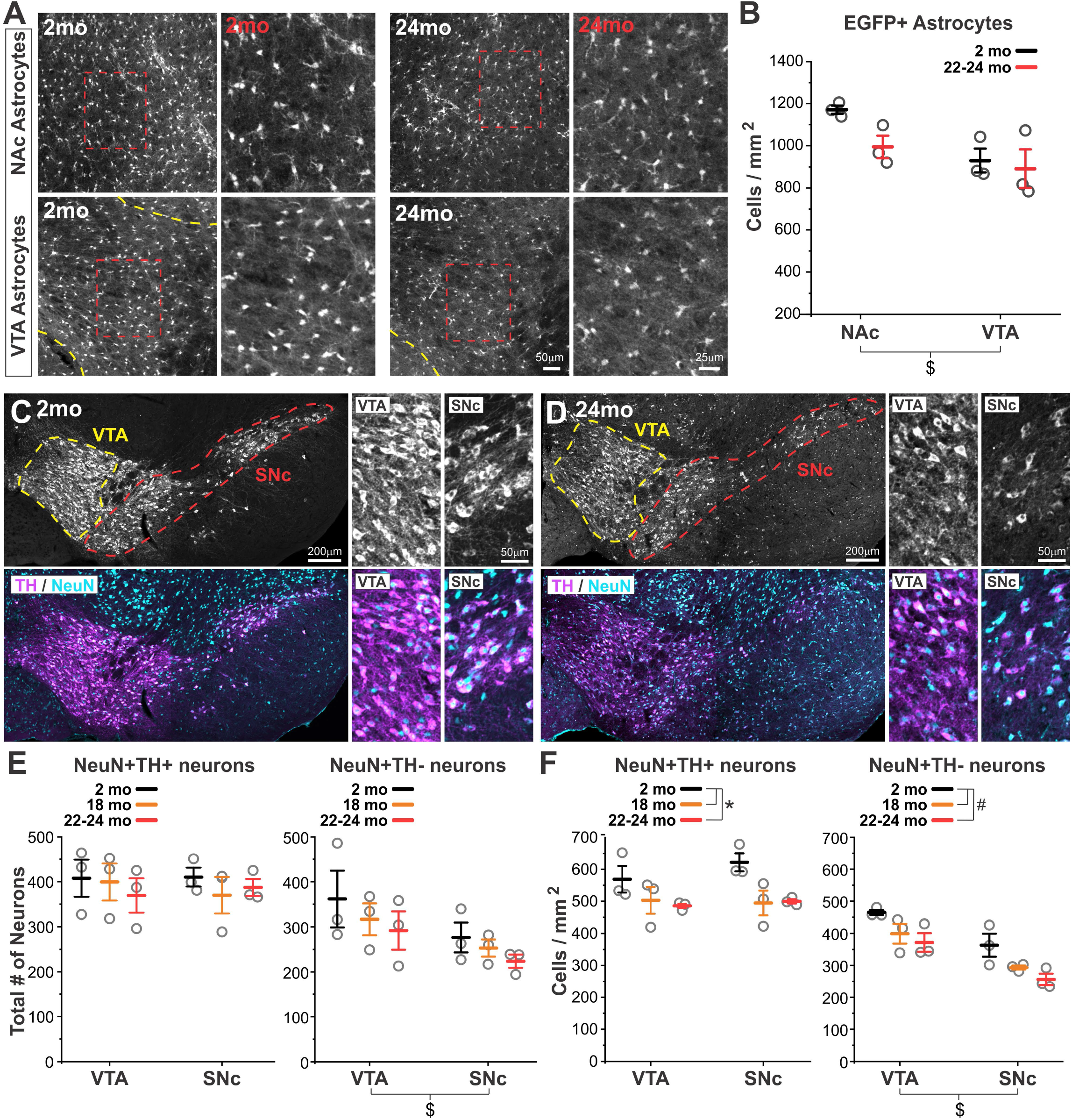
VTA and SNc microglial responses to aging are not accompanied by increases in local astrocyte abundance or significant neuronal loss. **A, B** – Images and quantification of NAc and VTA astrocyte density from 2 month (mo) and 24mo old *ALDH1L1-EGFP* mice. *Dashed yellow line* shows VTA boundary. Two-way ANOVA: main effect of age F(1,11) = 3.2, P = 0.11 n.s.; main effect of brain region F(1,11) = 8.1, P = 0.02; interaction F(1,11) = 1.3, P = 0.29 n.s. $ P = 0.02 NAc vs. VTA. N = 3 mice analyzed per age. **C, D** –Immunostaining for TH, which marks dopamine neurons, and NeuN which labels all neurons, in 2mo and 24mo *CX3CR1*^*EGFP /+*^ mice. VTA and SNc divisions shown with *yellow and red dashed lines*. Panels *at right* show higher magnification images of VTA and SNc neurons. Background autofluorescence (lipofuscin) is visible throughout 24mo tissue but does not interfere with clear identification of neurons. **E** – Total number of TH+ neurons and TH-negative (non-dopamine) neurons across 3 stereologically matched brain sections during aging. Two-way ANOVAS: **TH+ neuron sum** main effect of age F(2,17) = 43, P = 0.66 n.s.; main effect of brain region F(1,17) = 0.01, P = 0.92 n.s.; interaction F(2,17) = 0.24, P = 0.79 n.s.; **Non-TH neuron sum** main effect of age F(2,17) = 1.31, P = 0.30; main effect of brain region F(1,17) = 5.4, P = 0.04; interaction F(2,17) = 0.04, P = 0.96 n.s. $ P = 0.04 VTA vs. SNc. **F** – Density of TH+ neurons and TH-negative neurons during aging. Two-way ANOVAS: **TH+ neuron density** main effect of age F(2,17) = 6.8, P = 0.01; main effect of brain region F(1,17) = 0.51, P = 0.49 n.s.; interaction F(2,17) = 0.49, P = 0.63 n.s. * P < 0.03, 2mo vs. 18mo and 2mo vs. 22-24mo; **Non-TH neuron density** main effect of age F(2,17) = 8.97, P = 0.004; main effect of brain region F(1,17) = 29.64, P = 0.0002; interaction F(2,17) = 0.04, P = 0.96 n.s. # P < 0.05, 2mo vs. 18mo and 2mo vs. 22-24mo. $ P = 0.0002 VTA vs. SNc.

**Figure 8.**
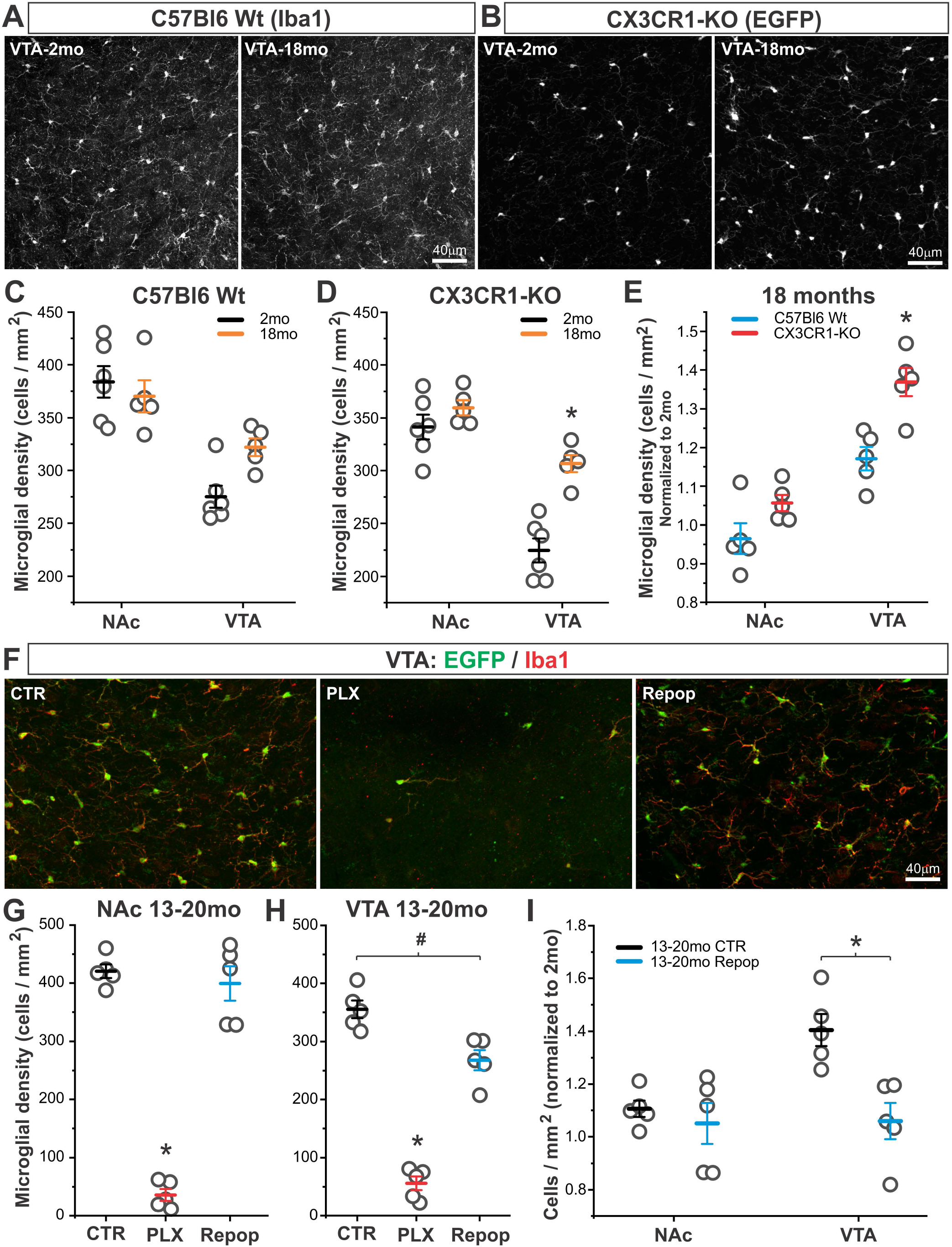
Region-specific microglial responses to aging are modulated by CX3CR1-signaling and microglial ablation/repopulation. **A, B** – VTA microglia of young and aging wildtype mice and *CX3CR1*^*EGFP /EGFP*^ (*CX3CR1-KO*) mice as revealed by Iba1 immunostaining and native EGFP, respectively. **C** – Microglial density in young and aging wildtype mice. Two-way ANOVA: main effect of age F(1,18) = 1.72, P = 0.21 n.s.; main effect of brain region F(1,18) = 37.9, P < 0.0001; interaction F(2,18) = 5.6, P = 0.03. **D** – Microglial density in young and aging *CX3CR1-KO* mice. Two-way ANOVA: main effect of age F(1,18) = 68.6, P < 0.0001; main effect of brain region F(1,18) = 23.9, P = 0.0002; interaction F(2,18) = 9.7, P = 0.006. * P = 0.0002, VTA 2 mo vs. VTA 18 mo. **E** – Microglial density when normalized to 2mo values from each brain region and genotype. Two-way ANOVA: main effect of genotype F(1,16) = 18.9, P = 0.0005; main effect of brain region F(1,16) = 63.4, P < 0.0001; interaction F(2,16) = 2.6, P = 0.12 n.s. * P = 0.004, VTA Wt vs. VTA KO. **F** –VTA microglia in 13-20mo *CX3CR1*^*EGFP /+*^ mice treated with control (*CTR*), Plexxikon 5266 CSFR1 inhibitor diet (*PLX*), or PLX followed by control diet for 21 days to allow for microglial repopulation (*Repop*). **G** – NAc microglial density. One-way ANOVA: F(2,14) = 125.4, P < 0.0001. * P < 0.0001, CTR vs. PLX and Repop vs PLX. **H** – VTA microglial density. One-way ANOVA: F(2,14) = 105.9, P < 0.0001. * P < 0.0001, CTR vs. PLX and Repop vs PLX. # P = 0.004, CTR vs. Repop. **I** – Microglial density when normalized to 2mo values from each brain region and treatment. Two-way ANOVA: main effect of treatment F(1,19) = 10.5, P = 0.005; main effect of brain region F(1,19) = 6.17, P = 0.02; interaction F(2,19) = 5.5, P = 0.03. * P < 0.007, VTA CTR vs. VTA repop.

## RESULTS

### VTA microglia show early increases in cell density during aging

Cortical and white matter microglia show significant increases in cell density as mice reach two years of age^6,17,21^. Whether increasing microglial abundance during aging is universal or varies across brain regions is largely unexplored. In young adult mice, microglia in distinct basal ganglia nuclei exhibit prominent regional differences in cell attributes, including cell density^13^. To determine whether these regional differences in cell density are maintained during aging, we analyzed microglia in the VTA and nucleus accumbens (NAc) of aging *CX3CR1*^*EGFP/+*^ mice.

In the NAc, microglial density remained largely consistent at 2 months (mo), 18mo, and 22-24mo (**Fig. 1A,B**). In contrast, in the VTA, microglial density increased significantly at 18mo, with trends toward further elevation at 22-24mo. These increases represent a roughly 28% increase in microglial abundance at 18mo and a 42% increase at 22-24mo. Moreover, at 2mo, microglial density was consistently higher in the NAc than the VTA of the same mouse (**Fig. 1C**). This inter-region relationship began to breakdown at 18mo and, by 22-24mo, VTA microglial density was similar to and sometimes exceeded that of microglia in the NAc of the same mouse. These findings reveal that expansion of the microglial population during aging is highly variable across distinct CNS regions.

### Resident microglia proliferate to drive expansion of the microglial population during aging

To determine the mechanisms underlying age-associated expansion of basal ganglia microglia, we first treated 18mo *CX3CR1*^*EGFP/+*^ old mice with BrdU. In both the NAc and VTA, morphological profiles of dividing cells as well as EGFP+BrdU+ microglia could be observed (**Fig. 2A,B**), suggesting that proliferation of resident microglia contributes to increases in microglial density during aging. Next, we used genetic approaches to tag microglia during young adulthood and to track their fate during aging. *CX3CR1*^*CreER-ires-EYFP*^ mice were crossed to mice expressing diphtheria toxin receptor preceded by a floxed-stopper module (*iDTR* mice). Double transgenic *CX3CR1*^*CreER-ires-EYFP*^*;iDTR* mice were injected with tamoxifen at 2mo of age and euthanized 2 weeks later or allowed to age to 20-24mo. Because mice do not normally express diphtheria toxin receptor, expression of this receptor acts as a “tag” to label microglia^18^ that were present in the CNS at 2mo of age (**Fig. 2C,D** and **Fig. S1**).

In mice euthanized 2 weeks after tamoxifen injection, 95% of Iba1+ or EYFP+ microglia were DTR+, demonstrating high recombination efficiency that allows tracking of the majority of microglia and any resulting progeny during aging (**Fig. 2E**). In mice that were euthanized at 20-24mo of age, overall microglial density had increased, consistent with observations in *CX3CR1*^*EGFP/+*^ mice (**Fig. S1**). In 20-24mo old mice, 95% of Iba1+ or EYFP+ microglia were also DTR+ (**Fig. 2E**). Maintenance of this percentage of DTR+ microglia in aged tissue would only be possible if microglia tagged at 2mo of age played a dominant role in proliferating and expanding the cell population. Indeed, the density of DTR+ microglia increased during aging and DTR+ cells exhibiting morphological profiles of cell division were evident in aged mice (**Fig. S1**). Furthermore, the fold change increase in DTR+ microglia was almost identical to the fold change increase in density of the total microglial population (**Fig. 2F**). While these analyses do not rule out the possibility that peripheral cells can enter the CNS during aging and adopt microglial phenotypes^26,27^, they indicate that the majority of observed microglial population expansion is driven by proliferation of CNS resident microglia.

It is worth noting that we observed a significant increase in NAc microglial density in these mice by 20-24mo of age, which we did not observe in the *CX3CR1*^*EGFP/+*^ mice. The *CX3CR1*^*CreER-ires-EYFP*^; *iDTR* mice and the *CX3CR1*^*EGFP/+*^ mice used in this study were aged in different vivariums and are on slightly different background strains (50% C57Bl6 50% BALB/cJ for *CX3CR1*^*CreER-ires-EYFP*^; *iDTR* mice and 100% C57Bl6 for the *CX3CR1*^*EGFP/+*^ mice). These observations raise the possibility that environment and genetic background may shape the overall timing and extent of microglial proliferation during aging.

### Morphological changes in basal ganglia microglia during aging

Cortical and hippocampal microglia exhibit altered morphology during aging and early stages of neurodegenerative disease^28–30^. To define the impact of aging on basal ganglia microglial cell structure, we examined high magnification images from 18mo old *CX3CR1*^*EGFP/+*^ mice. In both the NAc and the VTA, the majority of microglia exhibited stellate, highly branched morphologies similar to those observed in young adult mice (**Fig. 3A**). To assess the overall spatial relationship between microglia and surrounding tissue, we quantified the extent to which microglial processes and somas covered the field of view. In the NAc, microglial tissue coverage increased from 46±1% in 2mo old mice to 52±2% in 18mo old mice. Similarly, in the VTA, tissue coverage increased from 27±1% at 2mo to 33±1% by 18mo (**Fig. 3B**), representing a roughly 1.13-1.2 fold increase in both regions. However, in the NAc, estimated tissue coverage of individual microglia tended to increase, whereas, in the VTA it tended to decrease (**Fig. 3C**). This difference is likely due to the fact that the number of microglia increases at 18mo in the VTA but not the NAc. Hence, individual NAc microglia may increase soma size, process thickness, or process branching to achieve an overall increase in tissue coverage by the NAc microglial population. In the VTA, despite increased microglial density and increased population tissue coverage at 18mo, regions of “open space” devoid of microglial processes also appear increased (**Fig. 3A**). This observation would be consistent with a reduced process branching and territory of individual VTA microglia (**Fig. 3C**).

Some microglia in 18mo old mice also displayed morphological abnormalities in the form of somatic protrusions (**Fig. 3D**) resembling the beginnings of cell division. However, these protrusions did not contain DAPI+ nuclei and, instead were filled with CD68+ lysosome material. These structures were observed in roughly 3.4±2% of NAc microglia and 5.6±2% of VTA microglia. Collectively, these findings indicate that subtle changes in microglial morphology are detectable by late middle age and that such changes vary across brain region.

### Local increases in inflammation parallel expansion of the VTA microglial population

To determine the relationship between microglial population expansion and aging-induced CNS inflammation, we microdissected NAc and VTA tissue from 2mo, 18mo, and 22-24mo old *CX3CR1*^*EGFP/+*^ mice, and used qPCR to examine levels of key inflammatory signaling factors (**Fig. 4A-C**). We also analyzed microdissected mPFC, which is known to show increased inflammation during aging^31,32^. At 24mo of age, tumor necrosis factor alpha (TNFα) and interleukin 1 beta (IL-1β) showed a trend toward increased levels of expression within mPFC, and in the NAc and VTA, these increases were significant. Moreover, increases in TNFα and IL-1β were already significant at 18mo of age within the VTA (**Fig. 4B**). These regional differences in expression of inflammatory factors were particularly evident when comparing expression across regions within individual mice (**Fig. 4C**). At 2mo of age, levels of TNFα and IL-1β were consistently greater in the mPFC than in the NAc or VTA. Instead, at 22-24mo of age, levels of TNFα and IL-1β in the VTA were consistently greater than those observed in the NAc and mPFC of the same mouse.

To examine local inflammation at the protein level, we microdissected NAc and VTA tissue from 2mo, 18mo, and 22-24mo old *CX3CR1*^*EGFP/+*^ mice and used high sensitivity ELISA arrays to examine a panel of pro- and anti-inflammatory cytokines. Multiple pro- and anti-inflammatory cytokines were found at higher concentrations in the VTA compared to the NAc at all ages examined (**Fig. 4D** and **Fig. S2**). These included TNFα, IL-1β, interleukin 6 (IL-6), interferon gamma (IFNγ), interleukin 10 (IL-10), and interleukin 5 (IL-5). In the VTA, TNFα was further modulated by age, showing significant increases in tissue concentration at 22-24mo relative to 2mo. Some factors, such as chemokine ligand 1 (CXCL1; KC/GRO), were found at similar levels across both regions and were not modulated by age. Collectively these analyses suggest that inflammatory signaling is elevated locally within VTA and that this local elevation is further exacerbated during aging.

### Aging-induced changes in VTA microglia emerge by 13 months of age

To gain further insight into how early in the lifespan changes in VTA microglial phenotype emerge, we quantified microglial density and inflammatory factor production in 13-14mo old mice. In 13mo old *CX3CR1*^*EGFP/+*^ mice, significant increases in microglial density were detected within the VTA, but not the NAc (**Fig. 5A,B**). To examine local inflammatory factors with cellular specificity, we microdissected larger portions of midbrain, striatum, and cortex to allow FACS isolation of microglia and subsequent analysis of gene expression via digital droplet PCR (**Fig. 5C-E**). Microglial expression of TNFα tended to be greater in the midbrain compared to the striatum and cortex at all ages examined. Within individual mice, no significant differences across region were observed in 2mo old mice (**Fig. 5E**). In contrast, in 14mo old mice, midbrain microglia expressed significantly higher levels of TNFα transcript than striatal or cortical microglia in the same mouse. These analyses indicate that region-specific changes in microglial phenotype begin midway through the lifespan for mice, suggesting that chronic exposure to these “aged” microglia and a low-level inflammatory milieu through the remaining lifespan could critically shape synaptic function and disease vulnerability of nearby neurons.

### Similar aging-induced changes are evident in SNc microglia

Midbrain dopamine neurons are particularly vulnerable to functional decline and neurodegenerative disease during aging^33,34^. The underlying causes of this vulnerability are still poorly understood. Although dopamine neurons in the VTA exhibit elevated disease susceptibility, dopamine neurons in the SNc are typically more vulnerable to damage and decline^34^. To explore whether similar, aging-induced changes in microglial phenotype are observed in the SNc, we examined microglial density and morphology within the SNc of *CX3CR1*^*EGFP/+*^ mice. Microglial density in the SNc was also significantly elevated at 18mo and remained elevated at 22-24mo (**Fig. 6A** and **Fig. S3**). When normalized to densities in 2mo old mice, the increases in microglial density in the SNc were of similar magnitude to those in the VTA (**Fig. 6B**).

Similar to observations in NAc and VTA, morphological analysis revealed that there was also a roughly 1.2-fold increase in microglial tissue coverage by 18mo in the SNc (**Fig. 6C** and **Fig. S3**). Morphological abnormalities were also evident in the SNc (**Fig. 6D** and **Fig. S3**). Frequency of microglial somas with large protrusions was roughly 5.7±2%, similar to that observed in VTA (5.6±2%). There were also instances of large inclusions along cell processes that did not contain nuclear material and were usually filled with CD68. Qualitatively, these inclusions did not fully resemble phagocytic cups, which tend to appear more hollow and show a brilliant rim of EGFP^35^. These process protrusions were observed in roughly 4.5±2% of microglia in the NAc, 6.1±4% of microglia in the VTA, and 8.0±3% of microglia in the SNc.

One morphological abnormality appeared to be largely unique to the SNc: apparent “clusters” of microglial cells adjacent to or among SNc dopamine neurons (**Fig. 6E** and **Fig. S3**). Microglial clusters were observed in the SNc of most mice, were only rarely observed in the VTA and were not observed in the NAc (**Fig. 6F**). Although these clusters appeared to contain many microglial cells, co-labeling with DAPI revealed that they typically only contained 3-4 microglia that had an elaborated, extensive membrane surface (**Fig. 6G** and **Fig. S3**). No pyknotic nuclei indicative of dying cells were observed within these structures but they did contain abundant CD68+ lysosomes, suggesting degradation of engulfed material. Collectively, these analyses indicate that changes in the SNc microglial population during aging are largely similar to those observed in VTA microglia.

### Aging induced changes in VTA/SNc microglia precede systemic inflammation, local changes in astrocyte abundance and parallel early loss of neurons

Systemic inflammation increases with aging and regional differences in CNS penetration of peripheral cytokines^36^ could potentially contribute to early changes in VTA/SNc microglial phenotype. To explore this possibility, we measured serum levels of inflammatory signaling factors using high sensitivity ELISA cytokine arrays in 2, 18, and 22-24mo old *CX3CR1*^*EGFP/+*^ mice. Multiple pro-inflammatory factors, such as IFNγ, IL-6, and TNFα were significantly increased during aging (**Fig. S4**). Several factors that have a mix of pro- and anti-inflammatory functions or largely anti-inflammatory functions were also elevated during aging, including IL-10, IL-2, and CXCL1. Other factors such as IL-1β, IL-4, and IL-5 were detected but did not show significant changes across the ages sampled. In all cases where serum cytokines increased during aging, posthoc analyses revealed that these increases did not reach significance until 24mo. These data suggest that regional differences in CNS penetration of peripheral cytokines are unlikely to underlie early changes in VTA/SNc microglial phenotype and population expansion observed at 13-18mo of age.

CNS astrocytes make critical contributions to tissue homeostasis, produce inflammatory factors, and proliferate in response to CNS perturbations^28^. Microglia and astrocytes can regulate one another’s attributes^37^ and local abundance of basal ganglia microglia is tightly correlated with astrocyte density in young adult mice^13^. To explore whether regional differences in astrocyte responses to aging regulate regional differences in microglial aging, we examined astrocytes in young adult and aging *ALDH1L1-EGFP* mice. Astrocyte cell density remained relatively stable in the NAc and VTA between 2mo and 24mo of age (**Fig. 7A-B**), with a trend toward decreased astrocyte abundance in the NAc at 24mo. High magnification images from aging *ALDH1L1-EGFP* mice revealed that, qualitatively, astrocytes maintain their morphological characteristics, including interaction with local blood vessels (**Fig. S5**) and do not exhibit somatic protrusions or cell process inclusions like those observed in microglia. Together, these observations indicate that increases in VTA/SNc microglial density during aging are not caused by parallel increases in astrocyte abundance.

Microglia respond to death of nearby neurons with numerous changes in cell attributes. Because dopamine neurons are more susceptible to functional decline, early loss of these vulnerable neurons could potentially underlie early responses of VTA/SNc microglia to aging. To explore this possibility, we used two approaches to quantify the abundance of dopamine neurons in the VTA and SNc at 2, 18, and 24mo in *CX3CR1*^*EGFP/+*^ mice (**Fig. 7C,D**). Quantification of the total number of dopamine neurons in stereologically matched brain sections (**Fig. S5**) revealed trends toward reduced numbers of dopamine neurons at 18 and 24mo but these were not significant (**Fig. 7E**). Quantification of cell density revealed decreases in dopamine neuron cell density that were significant only when pooling data across the VTA and SNc (**Fig. 7F**), suggesting that loss of dopamine neurons is relatively minor at the ages examined. To check for cell death of glutamatergic and GABAergic neurons also present within the VTA and SNc ^34,38^, we quantified abundance of NeuN+TH-neurons. Similar to dopamine neurons, NeuN+TH-neurons showed trends toward reduced total number of neurons and significant reductions in cell density when pooling across the VTA and SNc (**Fig. 7E,F**). Collectively, these analyses suggest that neuron loss in the VTA/SNc is not extensive at the ages examined, in agreement with previous reports^39,40^, and that increases in VTA/SNc microglial proliferation and production of inflammatory factors likely precede rather than follow the beginnings of neuronal loss in these brain regions.

### CX3CR1 deletion and microglial ablation and repopulation modulate region-specific microglial aging responses

Signaling through CX3CR1 (fractalkine receptor) can modulate microglial properties^41,42^. Patterns of microglial gene expression associated with aging were more pronounced in mice lacking CX3CR1 and dopamine neuron loss more severe in MPTP-treated CX3CR1-lacking mice^43,44^. To determine whether signaling through this receptor could influence region-specific microglial responses to aging, we compared age-associated changes in microglial density in wildtype C57Bl6 mice and *CX3CR1*^*EGFP/EGFP*^ (*CX3CR1-KO*) mice (**Fig. 8A,B** and **Fig. S6**). Patterns of microglial abundance in these mice largely mirrored those observed in *CX3CR1*^*EGFP/+*^ mice. NAc microglia did not show significant changes in density between 2mo and 18mo of age in either C57Bl6 wildtype or *CX3CR1-KO* mice (**Fig. 8C,D**). VTA microglia instead showed trends toward significant increases in density in wildtype mice and significant increases in density by 18mo in *CX3CR1-KO* mice (**Fig. 8C,D**), highlighting consistency of this region-specific response of microglia to aging. Normalizing values to the mean density of microglia in each region at 2mo, revealed that the relative increase in VTA microglial density at 18mo was larger in *CX3CR1-KO* mice compared to wildtype mice. NAc microglial density in 18mo *CX3CR1-KO* mice also appeared slightly elevated compared to wildtype mice (**Fig. 8E**), leading to a relatively consistent ratio of VTA:NAc microglia within individual mice in both genotypes at 18mo (**Fig. S6**). These findings suggest that signaling through CX3CR1 can shape microglia responses to aging throughout the basal ganglia but that these effects are accentuated in regions prone to early microglial aging.

Microglia are dependent on activation of the colony stimulating factor 1 receptor (CSFR1) and can be ablated from the CNS via chronic pharmacological CSFR1antagonism^45^. Subsequent removal of CSFR1 antagonism allows microglia to repopulate the CNS^46^. Microglial ablation and repopulation in aging mice can partially reverse aging-induced changes observed in these cells and improve cognition^10,47^. To determine whether microglial ablation and repopulation can also mitigate early aging phenotypes of VTA microglia, we fed 13-20mo old *CX3CR1*^*EGFP/+*^ mice on PLX5622 or control diet for 21 days. PLX5622 resulted in an 91±0.2% and 84±1% reduction in NAc and VTA microglia, respectively (**Fig. 8F,G** and **Fig. S6**). Subsequent removal of PLX5622 for 21 days resulted in microglial repopulation in both regions. Within the NAc, microglial density in ablated/repopulated mice was comparable to that in control mice (**Fig. 8G**). Instead, in the VTA, microglial density in repopulated mice was significantly lower than that observed in control mice (**Fig. 8H**). Normalizing these values to the mean density observed in each region in 2mo old mice revealed that in control 13-20mo mice, VTA microglial density was roughly 1.4 fold elevated (**Fig. 8I**) but in 13-20 mo ablated/repopulated mice, it was equivalent to that observed in 2mo old mice. VTA microglial density was also consistently below that observed in the NAc of the same mouse following microglial ablation/repopulation (**Fig. S6**), similar to what is observed in 2mo old mice. Together, these results suggest that microglial ablation and repopulation during aging can, at least temporarily, mitigate region-specific expansion of the VTA microglial population.

## DISCUSSION

### Causes of microglial proliferation during aging

Here we demonstrate that the onset of microglial proliferation during aging varies widely and begins in midlife in select brain regions such as the VTA/SNc. Microglia are highly proliferative during early postnatal development as they colonize the CNS^26,35^ after which they reach a steady-state with minimal turnover in adulthood^48^. Nonetheless, numerous stimuli cause microglia to mount substantial proliferative responses throughout the lifespan including acute brain injury, disease processes, and repopulation after microglial ablation^29,30,46^. Studies disagree about the impact of aging on microglial abundance. Several studies report no change in substantia nigra^17^, progressive decreases in substantia nigra^49^, or transient decreases in VTA and SNc microglial number during aging^39^. However, exclusive reliance on Iba1 immunostaining, failure to distinguish substantia nigra *pars compacta* and *pars reticulata*, and use of 1 month timepoints as “young adult” comparisons complicates interpretation of these studies. Instead, others report that microglial number in cortex^6,50^, hippocampus^51^, and CNS white matter tracts^17,21^ increases late in life (22-24mo in mice or rats). We also observe increases in microglial abundance during aging, but find this can occur as early as middle age (13-14mo) (**Fig. 1**) in the VTA and SNc. The magnitude of microglial increases attained by late age in the VTA and SNc is much larger than that reported in cortex and on par with that reported in hippocampus and white matter. Moreover, by conditionally “tagging” microglia present in young adulthood via expression of DTR, we show for the first time that local expansion of the microglial population is chiefly driven by proliferation of resident CNS microglia (**Fig. 2 and Fig. S1**).

Causes of microglial proliferation during aging are less obvious than those in other contexts, such as populating the CNS during development, or clearing debris and dying cells after acute injury. Cell death seems unlikely to be the cause of microglial proliferation in the VTA/SNc during aging. We did not observe pyknotic nuclei, microglial phagocytotic cups, or substantial neuron loss in the VTA/SNc by 18mo of age (**Fig. 7**), despite prominent increases in microglial abundance at this time point. Similarly, inflammatory signaling also seems unlikely to be the primary cause of observed microglial proliferation. We did not detect increases in serum levels of inflammation until 24mo of age, and the local inflammation we observe within the VTA/SNc is likely milder than that observed during acute infection, injury, or degenerative disease^52^. Astrocytes also proliferate in response to infection-, injury-, or disease-induced inflammation^28^ and we did not observe significant increases in astrocyte number during aging. These observations suggest that chronic, low-level inflammation may be sufficient to raise vulnerability of local synapses and neurons but is likely not the major driver of microglial proliferation during healthy aging.

Future studies could explore the possibility that region-specific increases in microglial degradative burden or changing patterns of neuronal activity during aging underlie this early expansion of the VTA/SNc microglial population. Although we did not observe dying cells or microglial phagocytic cups, we did observe CD68+ microglial somatic protrusions and cell process inclusions in some cells and clusters of microglia within the SNc with abundant CD68. Microglia are equipped to sense and respond to changes in ongoing neuronal activity^53,54^ and increases in cortical microglial number during aging were modulated by loss of sensory input^50^, suggesting a role for neuronal activity in regulating microglial proliferation during aging. Behavioral studies in humans and rodents indicate that dopamine-dependent cognitive and motor performance declines during aging^40,55–60^, and excitability of dopamine neurons shifts during aging^61^, indicating that patterns of local neuronal activity are likely to be shifting within the VTA/SNc during aging and could potentially impact microglial proliferation.

### Chronic local inflammation as one consequence of microglial proliferation during aging?

During aging, increases in systemic and CNS inflammation have been observed across species^52,62,63^. We show that CNS inflammation during aging is not uniform; VTA (and likely SNc) neurons are exposed to chronic low-level inflammation throughout the lifespan with regional differences accentuating with age. Although we cannot unequivocally pinpoint the source of local inflammation, several observations suggest that microglia play a central role in regulating local inflammatory milieu during aging. First, the time course of VTA microglial proliferation is tightly aligned with the time course of increasing levels of TNFα and IL-1β transcript in this brain region. Furthermore, region-specific, age-associated increases in TNFα transcript were observed within midbrain microglia as early as 14mo of age. Finally, no parallel increases in astrocyte density were observed and published transcriptome data from aging astrocytes suggests that TNFα and IL-1β are not highly expressed by these cells in young or aged mice^64,65^. However, astrocyte expression of numerous genes does change during aging in a region-specific manner^64^, and our data do not preclude that astrocytes also shape how neuroprotective or neurotoxic the local environment is during aging.

The aging-induced increases in inflammation that we report are likely well below those experienced during active degenerative disease. Consistent with this idea, we observe relatively little VTA and SNc neuronal loss by 18 and 22-24mo of age. However, synapse loss and synapse dysfunction rather than overt neuronal death are most tightly linked to cognitive deficits during aging and presymptomatic stages of neurodegenerative disease^66–69^. Several observations suggest that perturbed synaptic function is a likely outcome of early, mild elevations in inflammation within the VTA/SNc. Age-dependent decline in dopamine-dependent cognition and movement is observed in humans^55–57,70^, non-human primates^71^, and rodents^40,60^, supporting the idea of perturbed synaptic function within this circuitry. The cytokines we find to be elevated within the VTA, TNFα and IL-1β, have both been shown to influence key aspects of glutamatergic and GABAergic synapse receptor composition, function, and placticity^72,73^. Finally, lifestyle manipulations that alter microglial phenotype and reduce inflammation, are correlated with improved cognition during aging^74,75^.

### How do region-specific responses of microglia to aging relate to disease susceptibility?

Although we do not observe significant dopamine neuron death during aging, several observations suggest that changing microglial attributes and chronic local inflammation in the VTA and SNc could contribute to increased disease susceptibility of dopamine neurons^33,34^. First, epidemiological studies indicate that long term use of non-steroidal anti-inflammatory medication reduced the risk of Alzheimer’s and Parkinson’s Disease (PD) by almost 50%^76–78^, suggesting that the local inflammatory milieu during aging does shape disease susceptibility. Second, accumulation of aggregate-prone proteins like α-synuclein is linked to risk of degenerative disease^79^. Although we did not directly assess microglial process motility or phagocytosis, the observation of altered microglial tissue coverage and increased “gaps” between microglial processes in VTA and SNc by 18 months suggests that microglial capacity to support aggregate clearance may be reduced. Finally, microglia respond to many of the environmental risk factors that increase PD susceptibility including traumatic brain injury and exposure to toxins^80–82^. Together these observations suggest that early changes in VTA and SNc microglia during aging lay a foundation for heightened vulnerability for nearby dopamine neurons that will intersect with other genetic and environmental risk factors to determine overall disease risk.

Two other factors that shape disease risk are worth exploring. First, men are at higher risk of developing PD than women^83^. Although male-female differences in microglial aging have been reported^84–86^, we did not observe any obvious male-female differences in the region-specific microglial changes that we report here (**Fig. S7**). Further research would be needed to unequivocally determine whether sex influences the degree of region-specific microglial responses to aging. Second, SNc dopamine neurons are more vulnerable to PD than VTA dopamine neurons, although both are eventually lost as disease progresses^33,34^. VTA and SNc microglia are much more similar to one another than they are to microglia in other basal ganglia nuclei or cortex^13^. VTA and SNc microglial responses to aging were also very similar in terms of increasing cell density, increases in tissue coverage, somatic CD68+ protrusions, and cell process inclusions. However, one morphological abnormality that was primarily observed in the SNc was the presence of microglial “clusters” among and adjacent to SNc dopamine neurons. These microglia clusters resemble “multinucleated giant cells (MGCs)” that have been reported in the context of presymptomatic amyotrophic lateral sclerosis^87^, HIV-induced dementia^88^, macaque and rat models of PD^63^, and in white matter tracts of aged (21 mo) mice^17^. Similar to our observations in the aging SNc, these studies report that MGCs rarely contained obvious pyknotic nuclei and that they tended to be arranged in circular fashion with nuclei around the periphery. There were no obvious differences between dopamine neurons near to these clusters and more distant from these clusters. *In vitro* experiments suggest that microglial MGCs are 2-4 fold more phagocytic and that their formation can be induced by stimulation with phagocytic challenges such as by amyloid-β and α-synuclein, as well as by inflammatory stimuli such as LPS, TNFα, and IFNγ^89^. While there is still little information available about microglial MGCs *in vivo*, their elevated presence in the SNc during aging suggests that this region is experiencing greater inflammatory signaling, phagocytic demand, and/or elevated α-synuclein than other nearby regions including the VTA. Additional research will be needed to determine if this is the case and if subtle differences in VTA and SNc microglial responses to aging further elevate the vulnerability of SNc dopamine neurons.

## Supporting information

Moca-Lecca Supplementary Figures

## Acknowledgments

This work was supported by the National Institute on Drug Abuse Intramural Research Program (L.M.D), the National Institute on Aging Intramural Research Program (N.G.), start-up funds from the David Geffen School of Medicine at UCLA (L.M.D.) and the Glen Foundation for Medical Research and American Federation for Aging Research Grant for Junior Faculty (L.M.D.). The authors thank Preethi Gopal, Chloe Noe, Sissi Palma Ribeiro, and Serena Scognamiglio for their contributions to early histology experiments and analysis. The authors thank the Johns Hopkins Bayview Immunomics Core (supported by NIAMS P30 AR-070254), Raffaello Cimbro, and the UCLA Eli and Edythe Broad Center of Regenerative Medicine and Stem Cell Research Flow Cytometry Core for assistance with FACS-based experiments. The authors thank the UCLA Eli and Edythe Broad Center of Regenerative Medicine and Stem Cell Research Microscopy Core for access to confocal microscopes. The authors thank Plexxikon for supplying the CSFR1 antagonist PLX5622. The authors thank Ari Schaler, Fanny Etienne, and Daniel Gray for help with manuscript editing.

## Author Contributions

Conceptualization, L.M.D; Methodology, L.M.D, E.N.M., D.L., D.T., N.H.G., and D.R.S.; Investigation: L.M.D, E.N.M., D.L., D.T., K.H., S.S., L.M., H.S., and R.M.; Formal Analysis, L.M.D, E.N.M., D.L., D.T., K.H., S.S., L.M., and H.S.; Visualization, L.M.D and E.N.M.; Writing, L.M.D.; Funding Acquisition, L.M.D, N.H.G, and D.R.S; Supervision, L.M.D, N.H.G, and D.R.S

