## Supplementary material for "Microglia drive pockets of neuroinflammation in middle age": Moca-Lecca Supplementary Figures

Supplementary Figure 1 - related to Figure 2

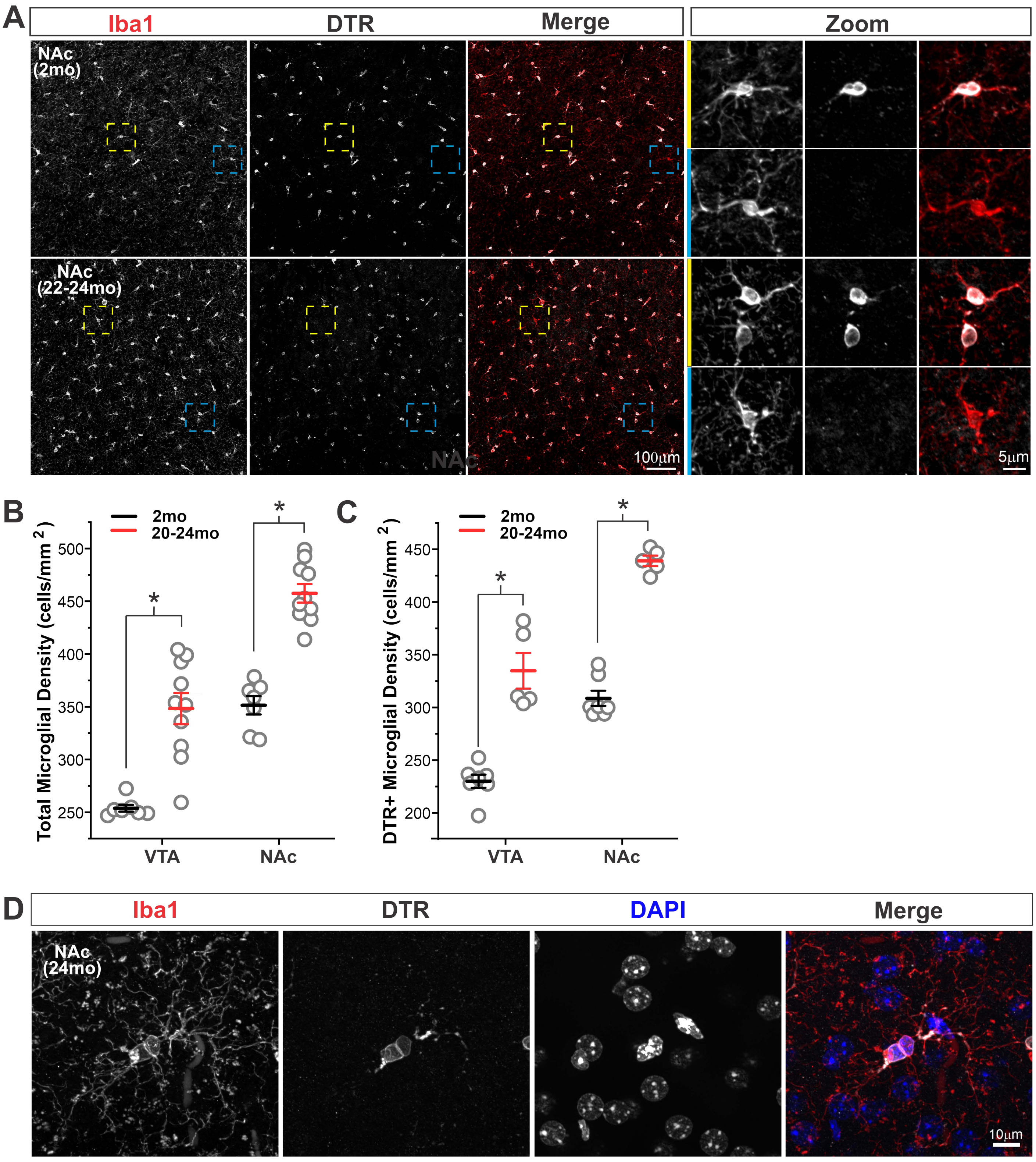

**Supplementary Figure 1. Microglia genetically tagged in young adulthood proliferate during aging. A –** Representative images from the NAc of 2mo and 20-24mo *CX3CR1*$^{CreER\text{-}IRES\text{-}EYFP}$*;iDTR* mice that were treated with tamoxifen at 2mo of age to label resident microglia present in the CNS at young adult ages. Examples of Iba1+DTR+ microglia (*yellow boxes*) and Iba1+DTR- microglia (*cyan boxes*) are shown at higher magnification in *panels at right*. **B –** Quantification of density of all microglia during aging in *CX3CR1*$^{CreER\text{-}IRES\text{-}EYFP}$*;iDTR* mice. Two-way ANOVA: main effect of age $F(1,23) = 160.3$, $P < 0.0001$; main effect of brain region $F(1,32) = 81.6$, $P < 0.0001$; interaction $F(1,32) = 0.2$, $P = 0.67$ n.s. * $P < 0.0001$.* P = **C –** Quantification of the density of DTR+ "tagged" microglia during aging in *CX3CR1*$^{CreER\text{-}IRES\text{-}EYFP}$*;iDTR* mice. Two-way ANOVA: main effect of age $F(1,32) = 78.8$, $P < 0.0001$; main effect of brain region $F(1,23) = 96.9$, $P < 0.0001$; interaction $F(1,23) = 1.9$, $P = 0.18$ n.s. * $P < 0.0001$. **D –** Representative example of DTR+ microglia exhibiting morphological profile of cell division.

### Supplementary Figure 2 - related to Figure 4

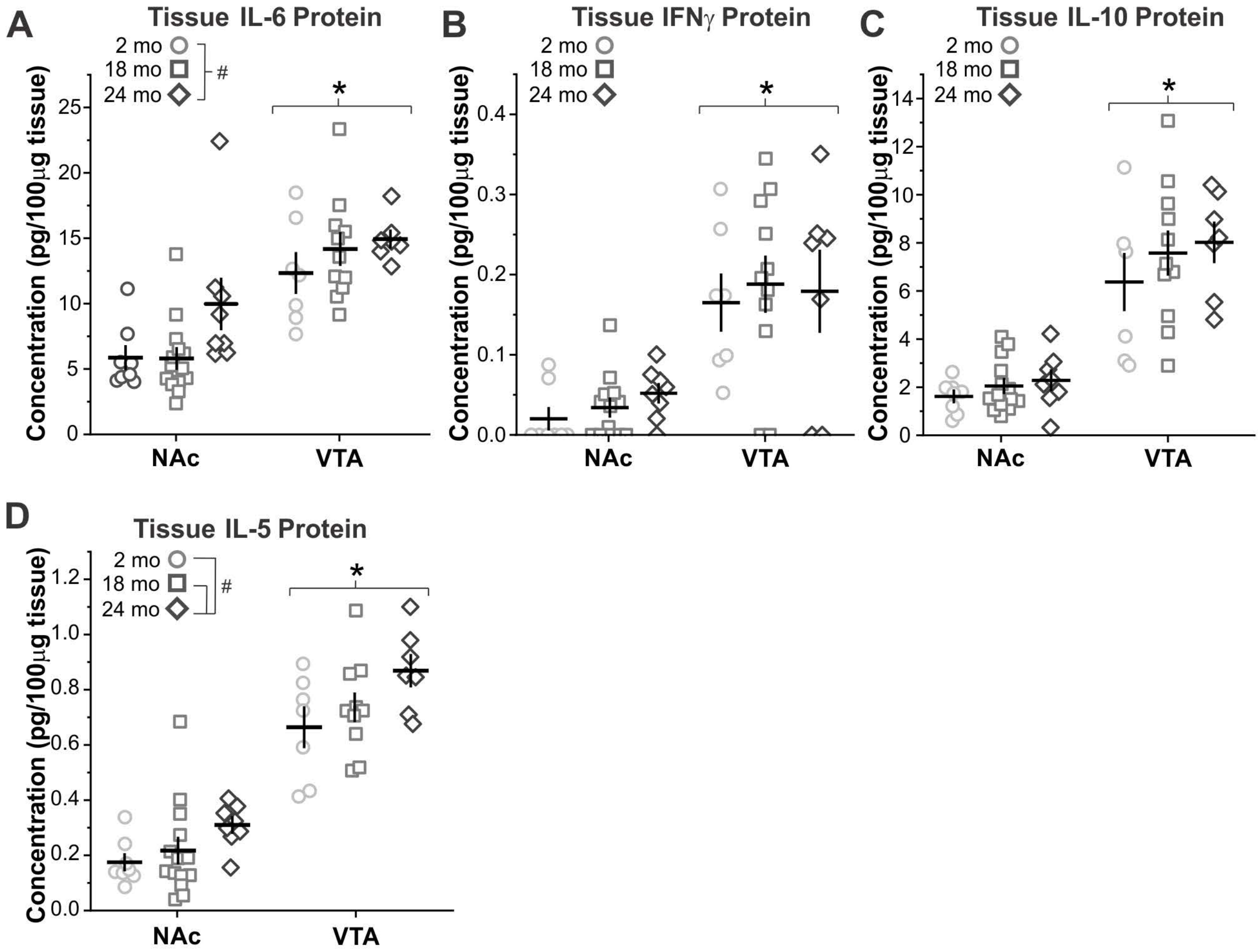

**Supplementary Figure 2. Tissue levels of multiple inflammatory factors are higher in the VTA than the NAc.** Tissue levels of inflammatory cytokines as measured by high sensitivity ELISA. **A** – IL-6 Two-way ANOVA: main effect of age $F(2,54) = 3.6$, $P = 0.03$; main effect of brain region $F(1,54) = 44.2$, $P < 0.0001$; interaction $F(2,54) = 1.1$, $P = 0.34$ n.s. * $P = 0.0001$, VTA vs. NAc. # $P = 0.04$, 24mo vs 2 mo. **B** – IFNγ Two-way ANOVA: main effect of age $F(2,53) = 0.35$, $P = 0.7$ n.s.; main effect of brain region $F(1,53) = 38.7$, $P < 0.0001$; interaction $F(2,53) = 0.12$, $P = 0.88$ n.s. * $P < 0.0001$, VTA vs. NAc. **C** – IL-10 Two-way ANOVA: main effect of age $F(2,54) = 1.3$, $P = 0.27$ n.s.; main effect of brain region $F(1,54) = 91.0$, $P < 0.0001$; interaction $F(2,54) = 0.25$, $P = 0.78$ n.s. * $P = 0.0001$, VTA vs. NAc. **D** – IL-5 Two-way ANOVA: main effect of age $F(2,54) = 5.2$, $P = 0.009$; main effect of brain region $F(1,54) = 161.0$, $P < 0.0001$; interaction $F(2,54) = 0.21$, $P = 0.81$ n.s. * $P = 0.0001$, VTA vs. NAc. # $P < 0.04$ 24mo vs. 2mo and 24mo vs. 18mo.

### Supplementary Figure 3 - related to Figure 6

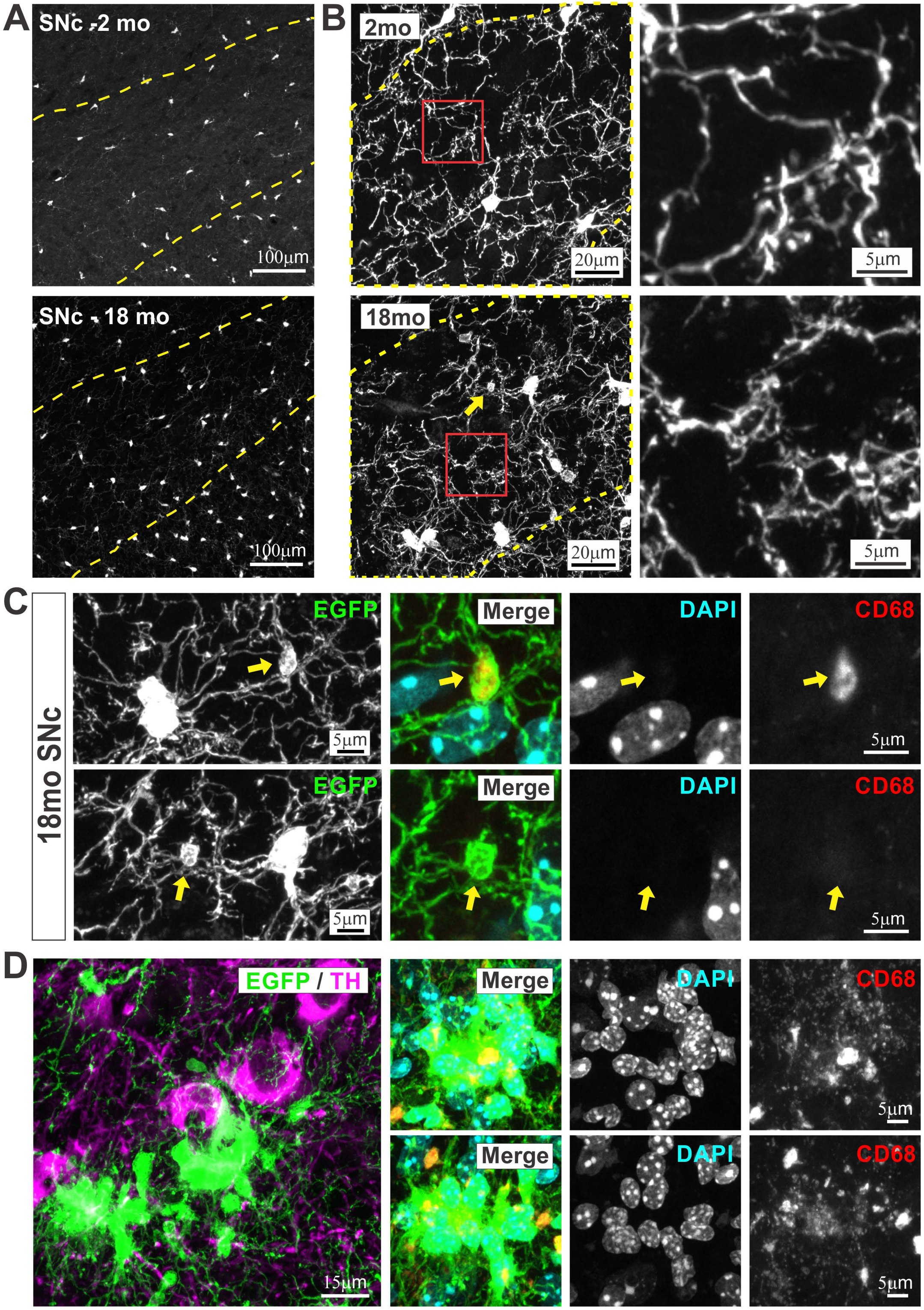

**Supplementary Figure 3. Changes in SNc microglia during aging parallel those observed in VTA microglia. A –** Representative images of microglia in the SNc of 2 month (mo) and 18mo old *CX3CR1*$^{EGFP/+}$ mice that were used for quantification of microglial density. *Dashed yellow line* represents the boundary of the SNc. **B –** Representative high magnification images of microglial branching structure in the SNc of 2mo and 18mo old *CX3CR1*$^{EGFP/+}$ mice used for quantification of microglial tissue coverage. Distal processes highlighted by *red boxes* shown at higher magnification in panels *at right*. *Yellow arrow* highlights an example of process swelling that is show at higher magnification in *C* (*lower panels*). **C –** High magnification examples of process swellings from SNc microglia in 18mo *CX3CR1*$^{EGFP/+}$ mice. *Yellow arrows* and co-staining with DAPI and CD68 indicate that these swellings do not represent instances of cell division or engulfment of dying cells with pyknotic nuclei and that these swellings frequently contain lysosomes (*top panels*), but occasionally do not (*bottom panels*). **D –** Additional examples of microglial cell clusters observed in the SNc of 18mo old *CX3CR1*$^{EGFP/+}$ mice. Co-staining of these clusters with DAPI and CD68 *at right* reveal that the clusters do not appear to contain engulfed pyknotic nuclei and that CD68 staining is abundant within the cluster, and appears to be composed of small punctate lysosomes more than the large 3-4μm CD68+ lysosomes observed in somatic and cell process swellings.

### Supplementary Figure 4 - related to Figure 7

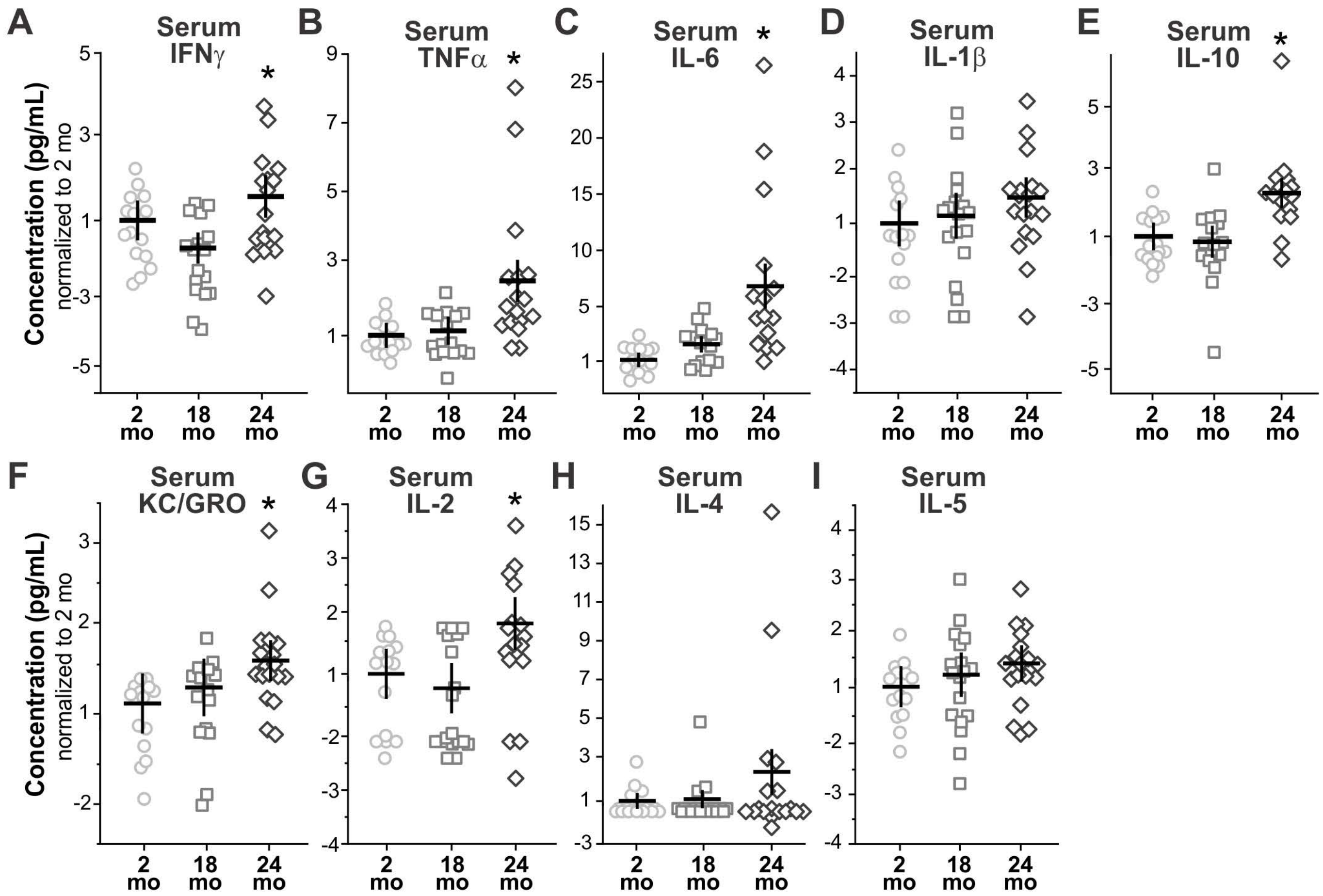

**Supplementary Figure 4. Systemic inflammation is not detectable in blood plasma until 24mo of age.** **A** – Serum interferon gamma (IFNγ) One-way ANOVA $F(2,49) = 5.28$, $P = 0.0085$; * $P = 0.007$, 24mo vs. 18mo. **B** – Serum tumor necrosis factor alpha (TNFα) One-way ANOVA $F(2,47) = 10.14$, $P = 0.0002$; * $P < 0.002$, 24mo vs. 18mo and 24mo vs. 2mo. **C** – Serum interleukin-6 (IL-6) One-way ANOVA $F(2,43) = 9.15$, $P = 0.0005$; * $P < 0.006$, 24mo vs. 18mo. and 24mo vs. 2mo. **D** – Serum interleukin-1 beta (IL-1β) One-way ANOVA $F(2,50) = 2.32$, $P = 0.11$ n.s. **E** – Serum interleukin-10 (IL-10) One-way ANOVA $F(2,48) = 10.16$, $P = 0.0002$; * $P < 0.001$, 24mo vs. 18mo. and 24mo. vs 2mo. **F** – Serum chemokine ligand 1 (CXCL1, KC/GRO) One-way ANOVA $F(2,47) = 4.23$, $P = 0.021$; * $P = 0.02$, 24mo vs. 2mo. **G** – Serum interleukin-2 (IL-2) One-way ANOVA $F(2,48) = 4.92$, $P = 0.011$; * $P < 0.009$, 24mo vs. 18mo. **H** – Serum interleukin-4 (IL-4) One-way ANOVA $F(2,50) = 1.44$, $P = 0.25$ n.s. **I** – Serum interleukin-5 (IL-5) One-way ANOVA $F(2,49) = 1.44$, $P = 0.25$ n.s.

Supplementary Figure 5 - related to figure 7

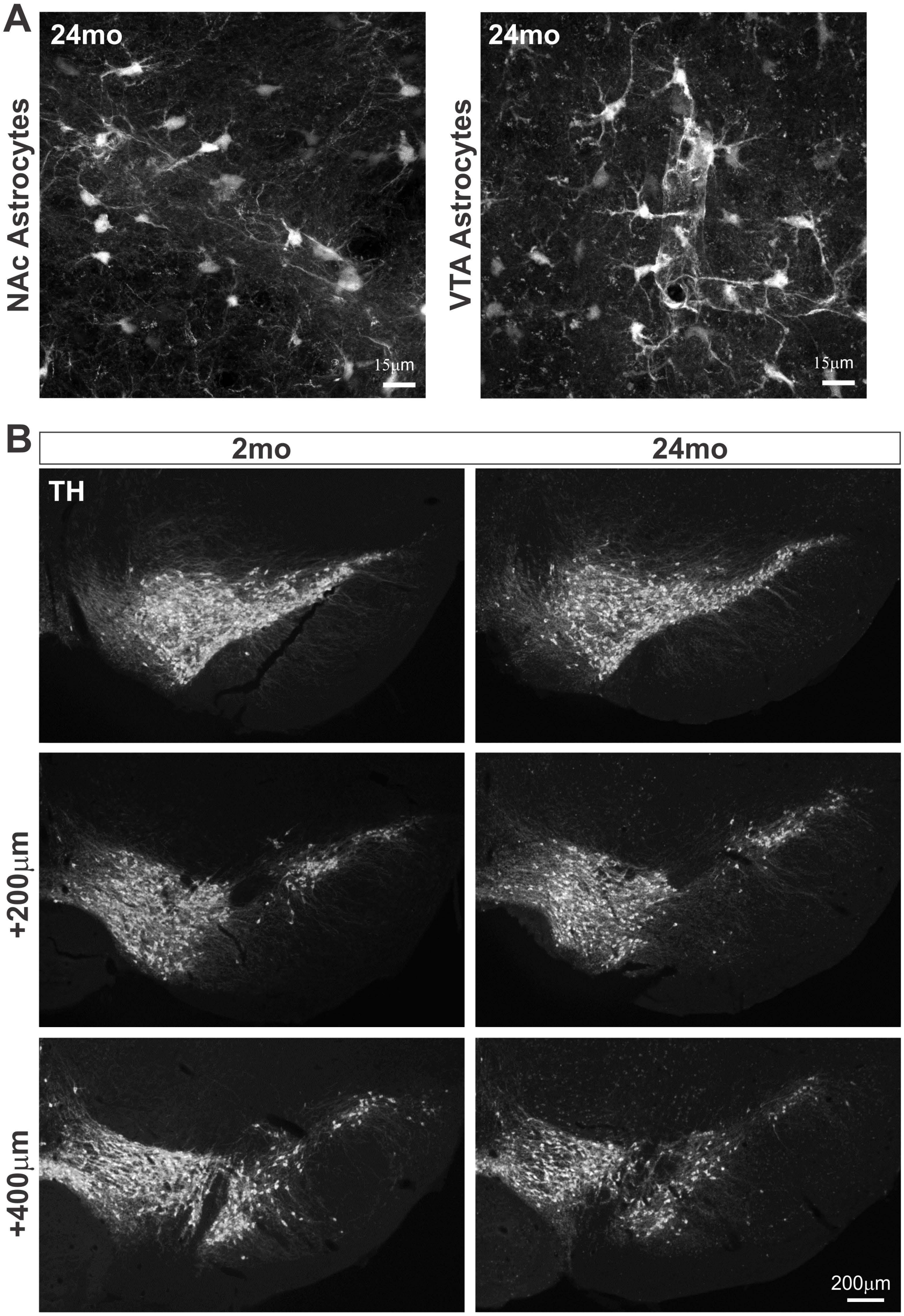

**Supplementary Figure 5. Astrocyte morphology in aging and stereological section sampling for neuron density quantification. A –** Sample high magnification confocal images from the NAc and VTA of 24mo *ALDH1L1-EGFP* mice showing ramified morphology and interaction with local vasculature similar to what is observed in young adult mice. **B –** Representative stereologically matched brain sections immunostained for tyrosine hydroxylase (TH) from 2 mo and 24 mo $CX3CR1^{EGFP/+}$ that were used for quantification of neuron abundance within the VTA and SNc.

### Supplementary Figure 6 - related to Figure 8

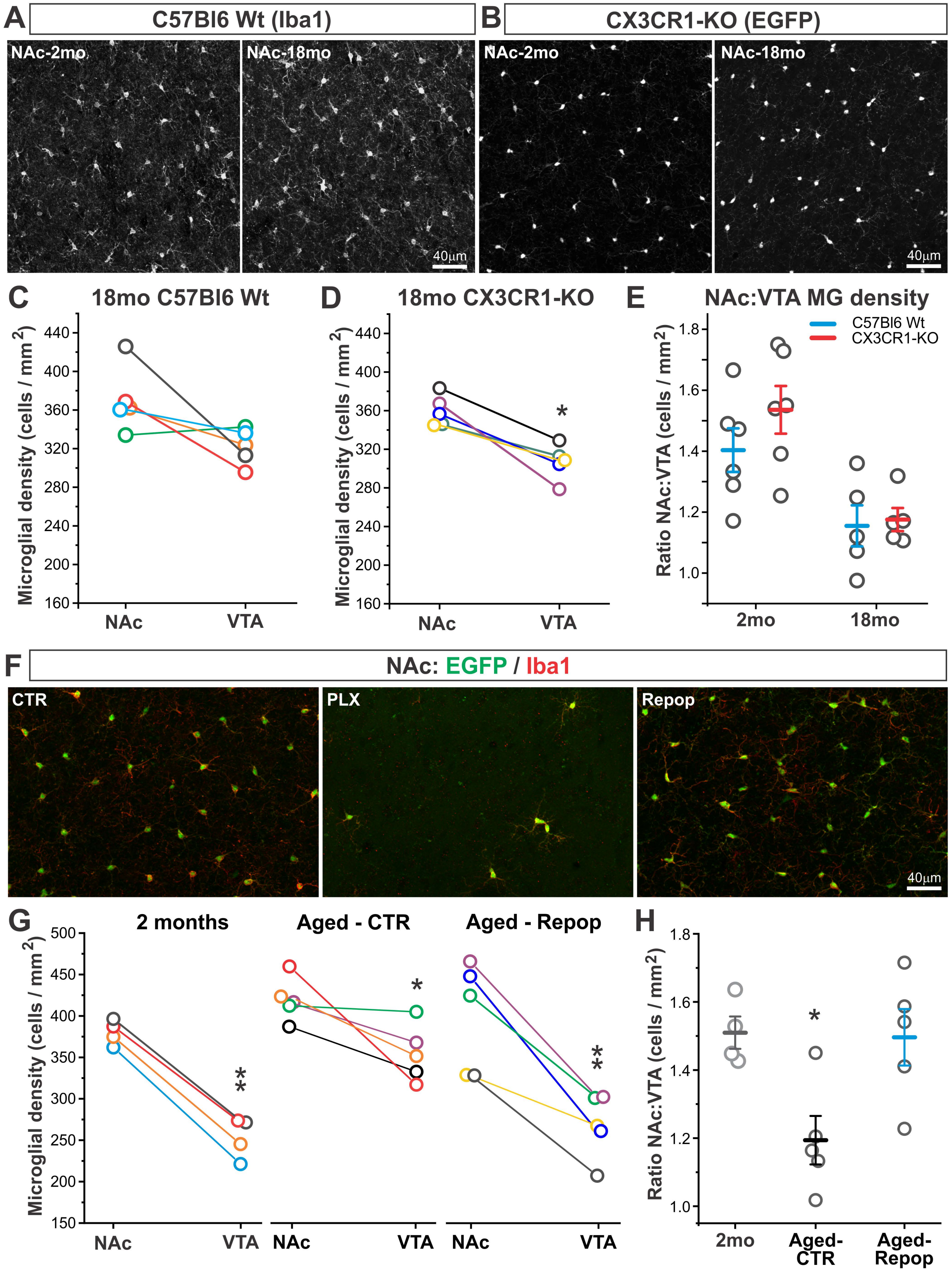

**Supplementary Figure 6. Region-specific microglial responses to aging are modulated by CX3CR1-signaling axis and microglial ablation/repopulation. A** – Representative confocal images of microglia in the NAc of young and aging wildtype mice as revealed by immunostaining for microglial marker Iba1. **B** – Representative confocal images of microglia in the NAc of young and aging *CX3CR1*$^{EGFP/EGFP}$ (*CX3CR1-KO*) mice. **C** – Comparison of NAc and VTA microglial density within individual young and aging wildtype mice. Data points from the same mouse are colored alike. P = 0.08, n.s., two-tailed paired t-test. **D** – Comparison of NAc and VTA microglial density within individual young and aging *CX3CR1-KO* mice. Data points from the same mouse are colored alike. * P = 0.006, two-tailed paired t-test. **E** – Ratio of NAc:VTA microglia within individual mice. Two-way ANOVA: main effect of age F(1,21) = 19.9, P = 0.0004; main effect of genotype F(1,21) = 1.3, P = 0.28 n.s.; interaction F(2,21) = 0.67, P = 0.42 n.s. **F** – Representative confocal images of NAc microglia in 13-20 mo *CX3CR1*$^{EGFP/+}$ mice treated with control (*CTR*) or Plexxikon 5622 CSFR1 inhibitor diet (*PLX*) as well as mice treated with PLX followed by control diet for 21 days to allow for microglial repopulation (*Repop*). **G** – Comparison of NAc and VTA microglial density within individual mice at 2 mo (same data as *Fig.1C*), 13-20 mo CTR mice, and 13-20 mo Repop mice. Data points from the same mouse are colored alike. At 2mo of age, VTA microglial density is less than NAc microglial density with high significance (** P = 0.0002, two tailed paired t-test). In 13-20 mo CTR mice, the relationship between NAc and VTA microglial density is still significant, but less so (*P = 0.04, two tailed paired t-test). In 13-20 mo Repop mice, VTA microglial density is again less than NAc microglial density with high significance (**P = 0.004, two tailed paired t-test). **H** – Ratio of NAc:VTA microglia within individual mice at 2 mo, 13-20 mo CTR, and 13-20 mo Repop. One-way ANOVA: F(2,13) = 105.9, P = 0.01. * P = 0.04, 2 mo vs. 13-20 mo CTR and 13-20 mo Repop vs. 13-20 mo CTR.

### Supplementary Figure 7 - Related to Figure 1 and 4

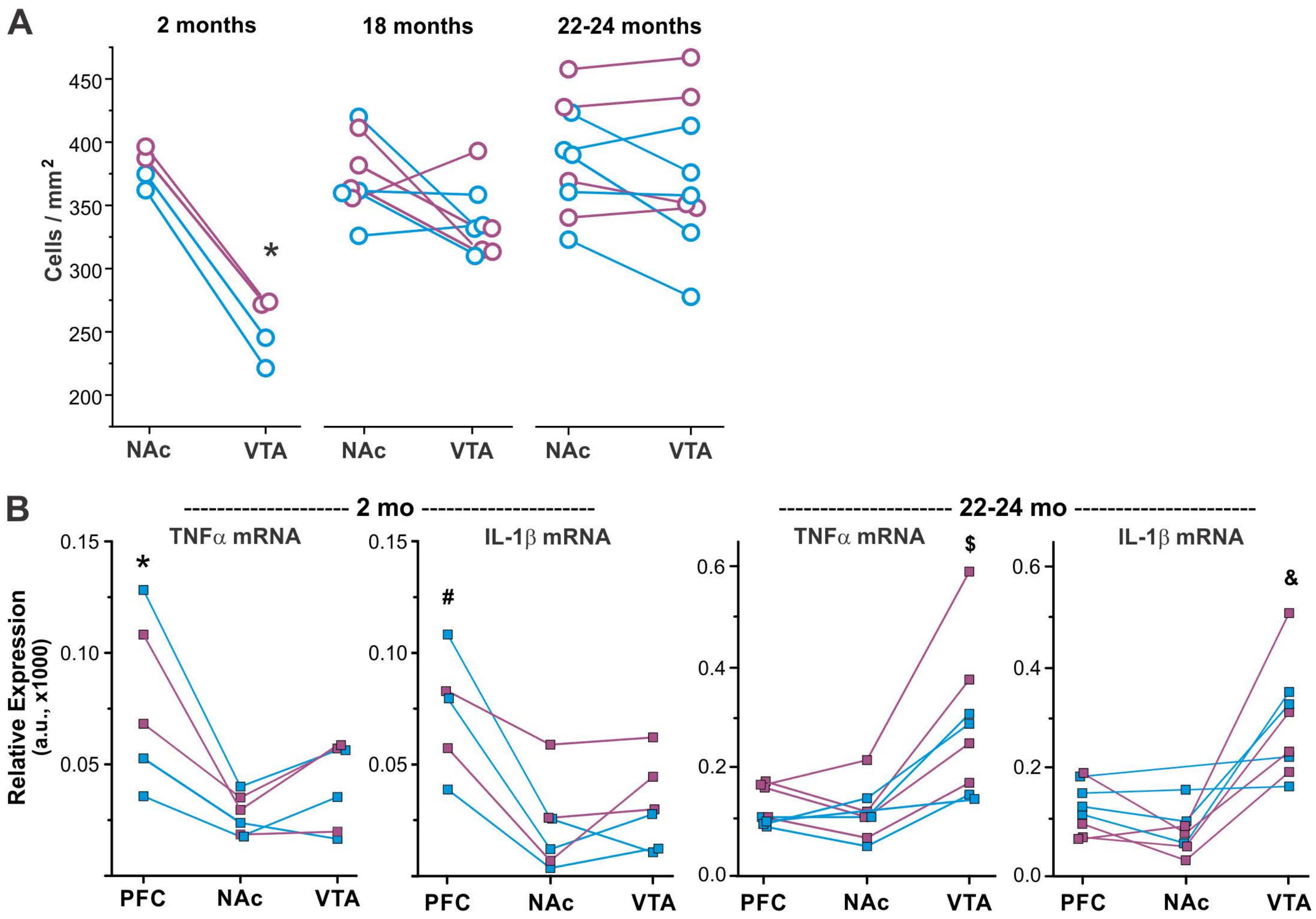

**Supplementary Figure 7. Prominent male-female differences are not observed in region specific responses of basal ganglia microglia to aging. A –** Graph of microglial cell density in the aging VTA and NAc from *Fig. 1C* with male mice colored *blue* and female mice colored *purple*. **B –** Graph of TNFα and IL-1β transcript in the VTA and NAc of 2mo and 22-24mo mice from *Fig. 4C* with male mice colored *blue* and female mice colored *purple*.
